# Test-retest reliability of regression dynamic causal modeling

**DOI:** 10.1101/2021.06.01.446526

**Authors:** Stefan Frässle, Klaas E. Stephan

## Abstract

Regression dynamic causal modeling (rDCM) is a novel and computationally highly efficient method for inferring effective connectivity at the whole-brain level. While face and construct validity of rDCM have already been demonstrated, here we assessed its test-retest reliability – a test-theoretical property of particular importance for clinical applications – together with group-level consistency of connection-specific estimates and consistency of whole-brain connectivity patterns over sessions. Using the Human Connectome Project (HCP) dataset for eight different paradigms (tasks and rest) and two different parcellation schemes, we found that rDCM provided highly consistent connectivity estimates at the group level across sessions. Second, while test-retest reliability was limited when averaging over all connections (range of mean ICC 0.24-0.42 over tasks), reliability increased with connection strength, with stronger connections showing good to excellent test-retest reliability. Third, whole-brain connectivity patterns by rDCM allowed for identifying individual participants with high (and in some cases perfect) accuracy. Comparing the test-retest reliability of rDCM connectivity estimates to measures of functional connectivity, rDCM performed favorably – particularly when focusing on strong connections. Generally, for all methods and metrics, task-based connectivity estimates showed greater reliability than those from the resting state. Our results underscore the potential of rDCM for human connectomics and clinical applications.

**Author Summary:** Test-retest reliability is an important prerequisite for the validity of connectivity estimates in many situations, particularly in clinical applications. Here, using different datasets from the Human Connectome Project, we demonstrate that regression DCM (rDCM) yields good to excellent test-retest reliability when focusing on strong connections. Comparing this to the test-retest reliability of functional connectivity measures, rDCM performed favorably in most cases. Furthermore, we show that reliability is not homogeneously distributed: we identified several regions (primarily in frontal and temporal lobe) that were linked via highly-reliable connections, regardless of the paradigm. Finally, we demonstrate that individual connectivity profiles are sufficiently unique that participants can be identified with high accuracy. Our findings emphasize the potential of rDCM for robust inference on directed “connectivity fingerprints” from fMRI data.

## 1 Introduction

Computational methods for assessing whole-brain effective (directed) connectivity from non-invasive neuroimaging data, such as functional magnetic resonance imaging (fMRI) or magneto-/electroencephalography (M/EEG) data, have great potential to further our understanding of brain function. This is because most, if not all, cognitive processes rests on widely distributed networks of neuronal populations (Bressler and Menon, 2010; McIntosh, 1999; Mesulam, 1990; Sporns, 2014). Similarly, directed connectivity measures at the whole-brain level are of major relevance for the young fields of Computational Psychiatry and Computational Neurology (Frässle et al., 2018b; Friston et al., 2014; Huys et al., 2016; Maia and Frank, 2011; Montague et al., 2012; Stephan and Mathys, 2014; Stephan et al., 2015). This is because dysconnectivity in large-scale networks has been postulated as a pathophysiological mechanism in various psychiatric and neurological disorders, such as, schizophrenia (Anticevic et al., 2015; Bullmore et al., 1997; Friston et al., 2016; Friston and Frith, 1995; Stephan et al., 2006), autism (Grèzes et al., 2009; Radulescu et al., 2013), major depression (Almeida et al., 2009; Schlösser et al., 2008; Vai et al., 2016), Parkinson’s disease (Dirkx et al., 2016; Marreiros et al., 2013), or epilepsy (Jirsa et al., 2016; Papadopoulou et al., 2017).

For clinical applications, computational methods for assessing functional integration in large-scale (whole-brain) networks of individual patients have great potential (Stephan et al., 2015). In order to leverage this potential, candidate methods need to fulfil several criteria, including (i) computational efficiency (allowing assessment of large-scale networks with hundreds of nodes, within clinically acceptable timeframes), (ii) reliability (construct and test-retest), and (iii) predictive validity (with regard to specific clinical questions).

Regression dynamic causal modeling (rDCM) is a generative model of fMRI data that was developed with these objectives in mind (Frässle et al., 2018a; 2017). It represents a novel variant of DCM for fMRI (Friston et al., 2003) which scales gracefully to very large networks including hundreds of nodes, enabling whole-brain effective connectivity analyses within timeframes of minutes to hours. Furthermore, the model can utilize structural connectivity information to constrain inference on directed functional interactions or, where no such information is available, infer optimally sparse representations of whole-brain connectivity patterns. For rDCM, we have recently demonstrated the face validity of the approach in comprehensive simulation studies both for task-based (Frässle et al., 2018a; 2017) and resting-state fMRI data (Frässle et al., 2021b). Furthermore, we have demonstrated its construct validity in application to fMRI data from a simple hand movement paradigm (Frässle et al., 2021c), as well as to resting-state fMRI data (Frässle et al., 2021b). These studies have provided promising results and suggest that rDCM might enable the construction of clinically useful “computational assay” in psychiatry and/or neurology (Stephan et al., 2015). However, test-retest reliability of rDCM has not been assessed so far.

Test-retest reliability represents an important test-theoretical property which quantifies the stability of estimates over time at the individual-subject level. It thus has particular relevance for clinical tests which require repeated assessments, such as monitoring treatment response over time. Test-retest reliability has already been assessed for classical variants of DCM for fMRI (Almgren et al., 2018; Frässle et al., 2016; 2015; Rowe et al., 2010; Schuyler et al., 2010). Overall, these studies have reported good reproducibility of DCM for fMRI across sessions, although detailed work has stressed avoidance of local extrema during optimization and the choice of the prior distributions as important factors for achieving good test-retest reliability (Frässle et al., 2015).

While test-retest reliability has been investigated for classical DCM for fMRI, it has not been tested for rDCM so far. Here, we assess the (group-level) consistency as well as the test-retest reliability of rDCM for inferring effective connectivity from task-based as well as resting-state fMRI data, applying the model to multiple datasets over time, acquired under the same conditions in the same participants. In addition, using the same data, we examined the consistency of group-level estimates of connectivity (referred to as “consistency” below). This metric is complementary to test-retest reliability which focuses on the stability of individual estimates over time. To this end, we made use of the comprehensive test-retest dataset from the Human Connectome Project (HCP; Van Essen et al., 2013).

## 2 Methods and Materials

### 2.1 Analysis plan

All analyses reported in this paper have been pre-specified in an analysis plan that was time-stamped prior to the analyses (https://gitlab.ethz.ch/tnu/analysis-plans/fraessle_hcp_test_retest).

### 2.2 Regression dynamic causal modeling

#### 2.2.1 General overview

Regression DCM (rDCM) is a novel variant of DCM for fMRI that enables effective connectivity analyses in whole-brain networks (Frässle et al., 2018a; 2017). This computational efficiency is achieved by several modifications and simplifications of the original DCM framework. These include (i) translating state and observation equations of a linear DCM from time to frequency domain, (ii) replacing the nonlinear hemodynamic model with a linear hemodynamic response function (HRF), (iii) applying a mean field approximation across regions (i.e., parameters targeting different regions are assumed to be independent), and (iv) specifying conjugate priors on neuronal (i.e., connectivity and driving input) parameters and noise precision. These modifications reformulate a linear DCM in the time domain as a Bayesian linear regression in the frequency domain, resulting in the following likelihood function:

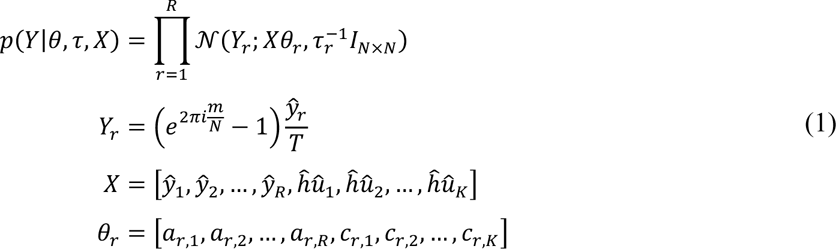

Here, *Y*_*r*_ is the dependent variable in region *r* that is explained as a linear mixture of afferent connections from other regions and direct (driving) inputs. Specifically, *Y*_*r*_ is the Fourier transformation of the temporal derivative of the measured signal in region *r*. Furthermore, *y*_*r*_represents the measured BOLD signal in region *r*, *X* is the design matrix (comprising a set of regressors and explanatory variables), *u*_*k*_ is the k^th^ experimental input, and the hat symbol denotes the discrete Fourier transform (DFT). Additionally, *θ*_*r*_ represents the parameter vector comprising all afferent connections *a*_*r*_, … , *a*_*r*_ and all driving input parameters *c*_r_, … , *c*_*r,k*_ targeting region *r*. Finally, *τ*_*r*_ denotes the noise precision parameter for region *r* and *I*_*N×N*_ is the identity matrix (where *N* denotes the number of data points). Choosing appropriate priors on the parameters and hyperparameters in Eq. (1) (see Frässle et al., 2017) results in a generative model that can be used for inference on the directed connection strengths and inputs.

Under this formulation, inference can be done very efficiently by (iteratively) executing a set of analytical Variational Bayes (VB) update equations of the sufficient statistics of the posterior density. In addition, one can derive an expression for the negative (variational) free energy (Friston et al., 2007). The negative free energy represents a lower-bound approximation to the log model evidence that accounts for both model accuracy and complexity. Hence, the negative free energy offers a sensible metric for scoring model goodness and is frequently used for comparing competing hypotheses (Bishop, 2006). We have recently further augmented rDCM by introducing sparsity constraints as feature selectors into the likelihood of the model in order to automatic prune fully connected network structures (Frässle et al., 2018a). A comprehensive description of the generative model of rDCM, including the mathematical details of the neuronal state equation, can be found elsewhere (Frässle et al., 2018a; 2017).

### 2.3 Dataset

#### 2.3.1 Participants

We used the publicly available fMRI data provided by the Human Connectome Project (HCP; Van Essen et al., 2013), specifically, all fMRI datasets from the HCP S1200 data release for which test and retest sessions are available. In total, this included 45 participants (31 females, 14 males). However, not all participants performed all paradigms twice. Hence, we excluded participants, for each paradigm individually, if not all their data from the test *and* retest session of the particular paradigm were available. The experimental protocol of the HCP was in compliance with the Declaration of Helsinki and was approved by the Institutional Review Board at Washington University in St. Louis (IRB #20120436). Informed consent was obtained from all participants prior to the experiment and all open access data were deidentified. Permission to use the open access data for the present study was obtained from the HCP, abiding the Data Use Terms (http://www.humanconnectome.org/data/data-use-terms).

#### 2.3.2 Data acquisition

The HCP dataset comprises fMRI data acquired during the “resting state” (i.e., unconstrained cognition in the absence of experimental manipulations). During the resting-state measurement, participants were asked to keep their eyes open and to fixate a crosshair projected on a screen. Furthermore, the HCP dataset comprises fMRI data acquired during several cognitive tasks, including: (i) working memory, (i) gambling, (iii) motor, (iv) language, (v) social cognition, (vi) relational processing, and (vii) emotional processing. For the resting state, a total of four measurements are available per session (i.e., test or retest) which differ in the phase encoding direction during oblique axial acquisitions. Specifically, two resting-state measurements are available with phase encoding in right-to-left (RL) and two in left-to-right (LR) direction. Similarly, for each task, two measurements are available (i.e., one per phase encoding direction) per session.

Functional images were acquired on the HCP’s custom 3T Siemens Skyra equipped with a 32-channel head coil. All fMRI data were acquired using a multiband accelerated 2D echo-planar imaging sequence (72 sagittal slices, TR = 720 ms, TE = 33 ms, voxel size 2 × 2 × 2 mm^3^, matrix = 104 × 90, flip angle = 52°, multi-band factor = 8) sensitive to the blood oxygen level dependent (BOLD) signal. Images covered the entire brain. The number of volumes acquired per measurement differed between paradigms: (i) resting-state = 1,200 volumes/measurement (approx. scan duration: 15 min), (ii) working memory = 405 volumes/measurement (5 min), (iii) gambling = 253 volumes/measurement (3 min), (iv) motor = 284 volumes/measurement (3 min), (v) language = 316 volumes/measurement (4 min), (vi) social cognition = 274 volumes/measurement (3 min), (vii) relational processing = 232 volumes/measurement (3 min), and (viii) emotional processing = 176 volumes/measurement (2 min).

For detailed information on the HCP dataset, please refer to the HCP S1200 manual (https://www.humanconnectome.org/storage/app/media/documentation/s1200/HCP_S1200_Release_Reference_Manual.pdf) or the relevant literature (Glasser et al., 2016b; Van Essen et al., 2013).

#### 2.3.3 Preprocessing

Preprocessing of the data was already performed by the HCP consortium and preprocessed files are released alongside the raw data. Here, we made use of the minimally preprocessed fMRI data (Glasser et al., 2013). The minimal preprocessing pipeline uses different tools from various freely available software packages like FSL (Jenkinson et al., 2012), FreeSurfer (Dale et al., 1999), and the HCP Workbench (Marcus et al., 2013) in order to accomplish several tasks, including spatial artifact/distortion removal, realignment, surface generation, cross-modal registration, and alignment to standard space (MNI). For the resting-state fMRI (rs-fMRI) data, additional preprocessing steps were performed to remove noise from the data. Specifically, the preprocessing of the rs-fMRI data made use of MELODIC as part of a single-subject spatial ICA decomposition. The resulting components were classified as signal or noise by FIX (Griffanti et al., 2014; Salimi-Khorshidi et al., 2014) and a cleaned version of the data is provided. The final preprocessed versions of both rs-fMRI and task data were then stored using the HCP-internal CIFTI file format and the associated grayordinates spatial coordinate system (Glasser et al., 2013). For comprehensive information on the individual preprocessing steps that were performed on both the HCP resting-state and task-based fMRI data, please refer to the manual (see above) or Glasser et al. (2013).

#### 2.3.4 Time series extraction

To extract BOLD signal time series for the subsequent rDCM analyses, we made use of two different whole-brain parcellation schemes. This allowed us to assess the robustness of our estimates of test-retest reliability and group-level consistency to the choice of parcellation scheme. First, we made use of the Human Connectome Project parcellation (HCP MMP 1.0; Glasser et al., 2016a), also known as the Glasser parcellation. HCP MMP 1.0 represents a very detailed cortical in vivo parcellation, consisting of 360 regions that were defined based on combined information on cortical architecture (e.g., relative cortical myelin content, cortical thickness), connectivity, and topography within some areas (e.g., the map of visual space in visual cortex). Second, we made use of the Schaefer 400 nodes parcellation (Schaefer et al., 2018), which rests on a gradient-weighted Markov Random Field model that integrates local gradient approaches (i.e., transient changes in functional connectivity patterns) and global similarity approaches (clustering of homogenous/similar functional connectivity patterns, regardless of spatial proximity). Using task-based and resting-state fMRI, the authors derive parcellations of the human brain at various degrees of granularity and demonstrate that these parcels represent subcomponents of global brain networks identified by Yeo et al. (2011). The Schaefer parcellation is optimized to align with both task-based and resting-state fMRI, and has been found to demonstrate improved homogeneity within parcels relative to alternative parcellations (Schaefer et al., 2018).

For each of the considered whole-brain parcellation schemes, we extracted the BOLD signal time series of all regions using dedicated HCP tools for CIFTI files. Specifically, we used the command *-cifti-parcellate* from the HCP Workbench tool *wb_command* (for further information, see https://www.humanconnectome.org/software/workbench-command/-cifti-parcellate). The script takes the dense time series data (which is the CIFTI format in which the HCP fMRI data are stored) and a *.dlabel file (which contains the parcellation) and extracts average BOLD signal time series from each region. The extracted time series then entered whole-brain effective connectivity analyses using rDCM.

#### 2.3.5 rDCM analysis

The extracted BOLD signal time series were then utilized for whole-brain effective connectivity analyses using rDCM. Since neither the Glasser atlas nor the Schaefer atlas are accompanied by an anatomical connectome that could inform the network architecture of the whole-brain models (i.e., the presence or absence of endogenous connections in rDCM; the A matrix), we assumed a fully (all-to-all) connected network. Furthermore, the input (C) matrix was defined according to data type: (i) for the resting-state fMRI datasets, no driving inputs are available and the C-matrix was set to all-zeros (as described in Frässle et al., 2021b), and (ii) for the task-based fMRI datasets, a full C-matrix was assumed.

For this setting, two different variants of rDCM were employed. First, using the fully connected network architecture, the strength of each connection and driving input was inferred using the classical implementation of rDCM (Frässle et al., 2017). This yielded a total of at least (i) 129,600 free parameters for the models based on the Glasser atlas (including 129,240 connectivity parameters, 360 inhibitory self-connections, and – for the task-based fMRI datasets – a task-dependent number of driving input parameters^1^), and (ii) 160,000 free parameters for the models based on the Schaefer atlas (including 160,000 connectivity parameters, 400 inhibitory self-connections, and – for the task-based fMRI datasets – a task-dependent number of driving input parameters).

In a second step, we utilized the sparsity constraints embedded in rDCM to automatically prune both connections and, for the task-based fMRI data, driving inputs (Frässle et al., 2018a). In brief, this is achieved by introducing binary indicator variables as feature selectors into the likelihood function where each indicator variable determines whether a specific connectivity parameter is present or not. This resulted in the same number of neuronal parameters (i.e., connectivity, inhibitory self-connection, and driving input parameters) as mentioned above, plus the same number of binary indicator parameters. Since exact *a priori* knowledge about the degree of sparseness of the networks is not available, we followed the procedure described in Frässle et al. (2018a) to determine the optimal *p*^i^_0_. More specifically, for each participant, we systematically varied *p*^i^_0_ within a range of 0.3 to 0.9 in steps of 0.1 and performed model inversion for each *p*^i^_0_ value. The optimal *p*^i^_0_ value was then determined for each participant by selecting the one that yielded the highest negative free energy. This yielded individual sparse effective connectivity patterns where some connections are absent (pruned away) and thus take a value of zero, whereas other connections remain present and thus take a non-zero connection strength.

For either of the two rDCM variants, the whole-brain models were fitted to the extracted BOLD signal time series by making use of the standard routines and prior settings implemented in the rDCM toolbox. Specifically, whole-brain models were inverted by utilizing the main routine *tapas_rdcm_estimate.m* from the rDCM toolbox as implemented in TAPAS (Frässle et al., 2021a), which is freely available as open source software (http://www.translationalneuromodeling.org/tapas).

#### 2.3.6 Group-level consistency and test-retest reliability of individual connection strengths

First, we investigated the across-session consistency of whole-brain effective connectivity patterns at the group level. To this end, for each endogenous connection and driving input, we computed the mean (across all participants) and then assessed the Pearson correlation between group-level parameter estimates from session 1 (“test”) and session 2 (“retest”). Significance was determined at an alpha-level of 0.05, corrected for multiple comparisons (i.e., number of paradigms) using Bonferroni correction. Hence, correlations with a p-value smaller than 0.00625 (i.e., 0.05/8) were deemed significant. These analysis steps were performed for both (i) rDCM with fixed network architecture, as well as (ii) rDCM with sparsity constraints.

Second, we assessed the test-retest reliability of the whole-brain effective connectivity patterns, i.e., the stability of rDCM parameter estimates at the individual-subject level. To this end, an intra-class correlation coefficient (ICC) was computed for each connection. Specifically, we utilized the ICC(3,1) type (Shrout and Fleiss, 1979), quantifying the ICC as a ratio between within-subject variability across the two sessions 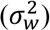 and between-subject variability 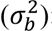:

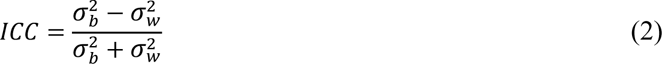

ICC(3,1) values range from -1 to 1. According to conventional interpretations of ICC values, test-retest reliability is classified as “poor” for ICC < 0.4, as “fair” for 0.4 ≤ ICC < 0.6, as “good” for 0.6 ≤ ICC < 0.75, and as “excellent” for ICC ≥ 0.75 (Cicchetti, 2001).

Based on the parameter-wise ICC values, different analyses were performed. First, the distribution of ICC values across all connections was inspected and the mean of the distribution was used to quantify the average test-retest reliability of rDCM when considering all connections. Second, reliability was tested as a function of connection strength. This was motivated by the hypothesis that reliability should be lower for connections that are weak (close to zero) and are thus unlikely to represent a meaningful effect that would be consistently present across sessions. Conversely, strong connections (both inhibitory and excitatory) should be more likely to represent meaningful effects and should thus have a greater probability to be conserved across sessions. This hypothesis was tested using two different analyses: (i) We computed the correlation between absolute parameter strengths and ICC values, and (ii) we restricted the test-retest reliability analyses only to parameters that were significantly different from zero (as assessed using one-sample *t*-tests and Bonferroni-correction for the multiple comparisons). Furthermore, for the connectivity parameters, we also further restricted the analysis to the top-1000 connections (i.e., the connections with the largest absolute weights).

#### 2.3.7 Inter-session consistency of whole-brain effective connectivity patterns

In a final analysis, we tested how consistent the entire effective connectivity profiles were across the two sessions. This analysis follows previous work demonstrating that individual subjects can be identified by their unique functional connectivity profiles derived from fMRI data (Finn et al., 2015). Here, we asked whether the whole-brain connectivity profile of individual participants during the first session (“test”) could be used to identify them from the set of all effective connectivity profiles obtained from the second session (“retest”). To this end, we computed for each participant in session 1 the similarity between his/her connectivity matrix and the connectivity matrices of all participants in session 2. The predicted identity was that with the highest similarity score. Following Finn et al. (2015), similarity was defined as the Pearson correlation between two vectors of connectivity estimates taken from the participant’s adjacency matrix from session 1 and all adjacency matrices from session 2. Repeating this procedure for each participant in session 1 allows us to construct a confusion matrix from which the identification accuracy can be computed. To account for order effects, we performed the same analysis in the opposite direction, testing whether a connectivity profile from the second session could be used to identify a given individual from the set of all effective connectivity profiles obtained from the first session.

To assess statistical significance of the identification accuracy, we performed permutation testing. Here, an empirical null distribution of the identification accuracy was computed by randomly permuting the participant labels of the session to be predicted and repeating the entire prediction procedure described above. Here, we used 1000 permutations. The p-value was then computed as the rank of the original identification accuracy in the distribution of permutation-based identification accuracies, divided by the total number of permutations.

## 3 Results

In the following, we first present our findings on group-level consistency and test-retest reliability of individual connection strength estimates. Subsequently, we report the inter-session consistency of whole-brain effective connectivity patterns. In either case, we present results obtained using both “classical” rDCM (with a fixed network architecture) and “sparse” rDCM (with sparsity constraints and thus variable network architecture). All results are compared to functional connectivity estimates (Pearson correlation coefficients and L1-regularized partial correlations).

### 3.1 Group-level consistency of connection strengths across sessions

#### 3.1.1 Regression DCM with fixed (fully connected) network architecture

Group-level estimates of individual connections were highly consistent across the two sessions, independently of the paradigm (i.e., task-fMRI, rs-fMRI) and whole-brain parcellation scheme. More specifically, for the Glasser atlas, Pearson correlations (r) for the connectivity parameter estimates ranged from 0.93 for the emotional processing task to 0.97 for the language task. For the driving input parameter estimates, Pearson correlations varied more strongly across the different paradigms and ranged from 0.37 for the emotional processing task to 0.98 for the social cognition task. For the Schaefer atlas, we found virtually identical results. A comprehensive list of all results from the group-level consistency analysis is provided in **Table 1**.

**Table 1.** Across-session consistency of group-level model parameter estimates for rDCM and functional connectivity. Consistency of parameter estimates in terms of the Pearson correlation coefficient between group-level (i.e., averaged across participants) estimates of session 1 (“test”) and session 2 (“retest”). Group-level consistencies are reported for the connectivity and driving input parameters of rDCM (*middle*) as well as for the functional connectivity estimates (*right*). For both methods, results are shown for all HCP paradigms as well as for the two whole-brain parcellation schemes (i.e., Glasser, Schaefer). Furthermore, results are reported for two different “modes” of estimation (see main text for details): (i) fixed network architecture (i.e., classical rDCM and Pearson correlation coefficient), and (ii) sparsity constraints (i.e., sparse rDCM and L1-regularized partial correlations). All correlations were significant at a significance threshold of *p* < 0.05 (Bonferroni-corrected for multiple comparisons).

|  | rDCM |  |  |  | FC |  |
| --- | --- | --- | --- | --- | --- | --- |
|  | *connectivity* |  | *inputs* |  |  |  |
| ***Fixed network*** |  |  |  |  |  |  |
|  | *Glasser* | *Schaefer* | *Glasser* | *Schaefer* | *Glasser* | *Schaefer* |
| REST | 0.96 | 0.96 | – | – | 0.95 | 0.94 |
| EMOTION | 0.92 | 0.91 | 0.37 | 0.38 | 0.89 | 0.87 |
| GAMBLING | 0.97 | 0.96 | 0.96 | 0.95 | 0.91 | 0.89 |
| LANGUAGE | 0.97 | 0.96 | 0.79 | 0.85 | 0.92 | 0.90 |
| MOTOR | 0.96 | 0.95 | 0.91 | 0.93 | 0.90 | 0.88 |
| RELATIONAL | 0.97 | 0.96 | 0.96 | 0.95 | 0.92 | 0.90 |
| SOCIAL | 0.97 | 0.97 | 0.98 | 0.97 | 0.92 | 0.90 |
| WORKING MEMORY | 0.95 | 0.95 | 0.89 | 0.90 | 0.90 | 0.88 |
| ***Sparsity constraints*** |  |  |  |  |  |  |
| REST | 0.61 | 0.62 | – | – | 0.98 | 0.98 |
| EMOTION | 0.90 | 0.89 | 0.66 | 0.61 | 0.91 | 0.91 |
| GAMBLING | 0.95 | 0.94 | 0.93 | 0.94 | 0.93 | 0.93 |
| LANGUAGE | 0.94 | 0.94 | 0.86 | 0.88 | 0.94 | 0.94 |
| MOTOR | 0.94 | 0.93 | 0.83 | 0.84 | 0.94 | 0.95 |
| RELATIONAL | 0.95 | 0.94 | 0.97 | 0.97 | 0.94 | 0.93 |
| SOCIAL | 0.95 | 0.95 | 0.97 | 0.97 | 0.95 | 0.95 |
| WORKING MEMORY | 0.90 | 0.90 | 0.92 | 0.92 | 0.95 | 0.95 |

#### 3.1.2 Regression DCM with sparsity constraints

In a second step, we assessed the across-session consistency of estimated connection strengths using rDCM with embedded sparsity constraints. Overall, we found group-level consistency of sparse rDCM to be comparable to rDCM with fixed network architecture for all paradigms except for the resting state. More specifically, for resting-state fMRI data, rDCM with sparsity constraints performed considerably worse (r = 0.62) than classical rDCM (r = 0.96); see **Table 1**. For all task-based datasets, consistency only slightly decreased for rDCM with sparsity constraints. Interestingly, for the driving input parameter estimates, rDCM with sparsity constraints performed comparable to rDCM with fixed network architecture and, in fact, in half of the cases outperformed the latter. For the Schaefer atlas, we again found results to be virtually identical.

#### 3.1.3 Comparison to functional connectivity

In a next step, we compared the group-level consistency of rDCM (both with fixed [fully connected] network architecture and sparsity constraints) with the group-level consistency of functional connectivity estimates that are frequently used for human connectomics and network neuroscience. Specifically, we assessed group-level consistency for functional connectivity estimates based on Pearson’s correlation coefficients (for a full connectivity matrix) and L1-regularized partial correlations (for sparsity constraints), respectively.

In brief, group-level Pearson correlations were highly consistent across the two sessions, regardless of the paradigm (i.e., task-fMRI, rs-fMRI) and whole-brain parcellation scheme. More specifically, for the Glasser atlas, group-level consistency for Pearson correlation coefficient ranged from 0.89 for the emotional processing task to 0.95 for the resting state (see **Table 1**). Hence, we found the group-level consistency for Pearson correlations to be somewhat lower than for rDCM. More specifically, we found differences to range between 0.01 and 0.06 (all in favor of rDCM), which was highly significant (*p* < 0.001) given the high degrees of freedom (i.e., number of connectivity parameters). For L1-regularized partial correlations, group-level consistency ranged from 0.91 for the emotional processing task to 0.98 for the resting state. Here, the values were generally very similar to sparse rDCM, except for the resting state dataset where L1-regularized partial correlations showed greater consistency. Except for the resting state, we found differences between sparse rDCM and L1-regularized partial correlations to range between 0.01 and 0.05 (in favor of one or the other), which was again highly significant (*p* < 0.001) given the high degrees of freedom. As for the rDCM analysis, we found functional connectivity results for the Schaefer atlas to be virtually identical to the ones for the Glasser atlas.

### 3.2 Test-retest reliability

#### 3.2.1 Regression DCM with fixed (fully connected) network architecture

In a second analysis, we assessed the test-retest reliability of estimates of individual connection strengths by rDCM, computing the ICC (Shrout and Fleiss, 1979) for each connection. Here, we report the results for the Glasser atlas; again, the results for the Schaefer atlas are virtually identical and are reported in the Supplementary Material.

Overall, when considering all model parameters, test-retest reliability of model parameter estimates from rDCM was relatively low (**Fig. 1B**, *left*). More specifically, for the connectivity parameters, on average test-retest reliability ranged from poor for the resting state (mean ICC = 0.24, 95% confidence interval (CI) = [-0.18 0.59]) to fair for the social cognition task (mean ICC = 0.42 [-0.07 0.75]) when considering all connections. Similarly, for the driving input parameters, test-retest reliability ranged from poor for the emotional processing task (mean ICC = 0.08 [-0.43 0.54]) to fair for the social cognition task (mean ICC = 0.42 [-0.03 0.73]). Importantly, this includes weak connections and driving inputs which may not represent meaningful effects, but may be driven by noise. In a next step, we therefore tested whether stronger parameters tended to be more reliable.

**Figure 1:**
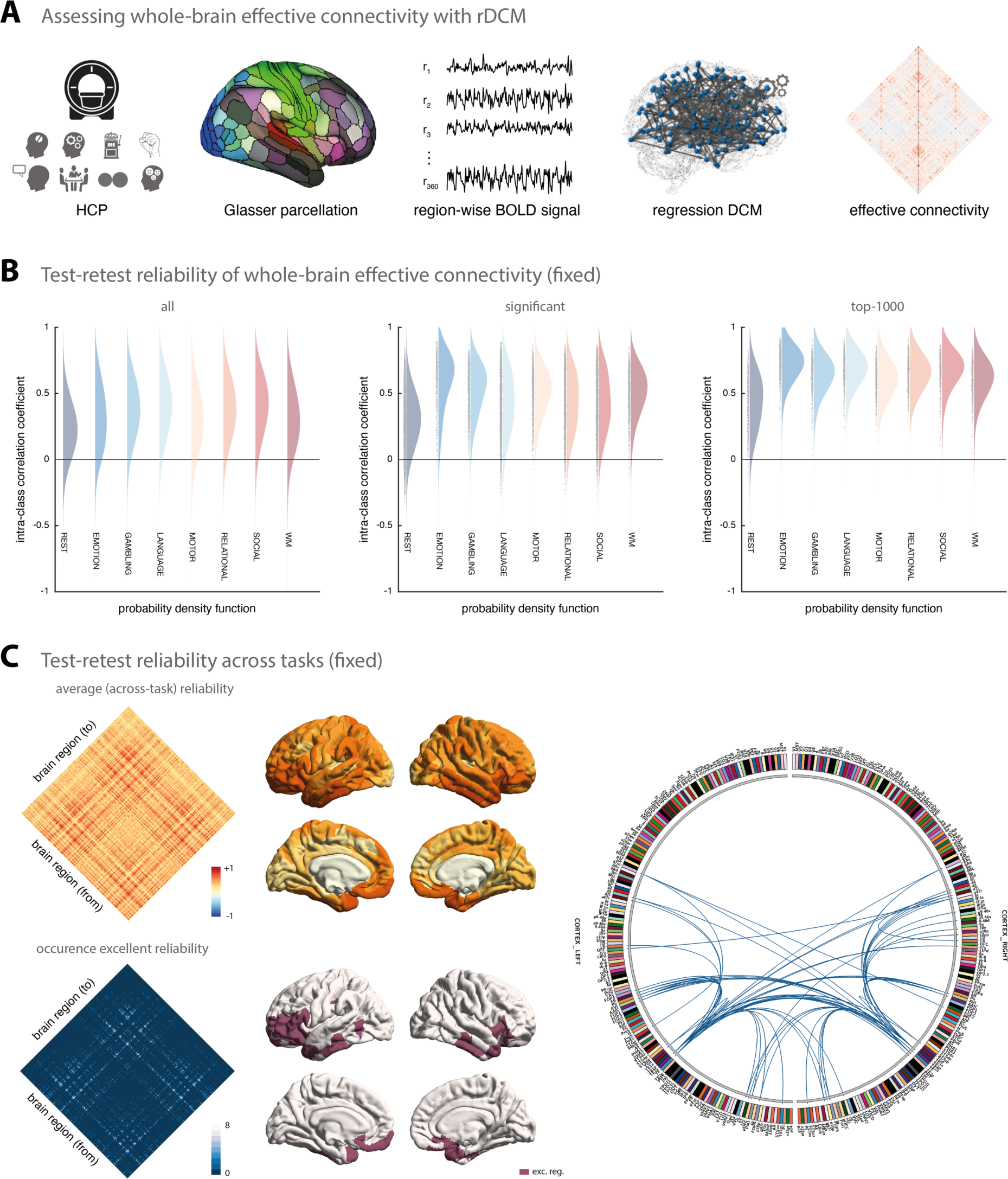
Test-retest reliability of regression DCM for a fixed network architecture. **(A)** Methodological overview. Resting-state and task-based fMRI data from the Human Connectome Project (HCP) is used for the analysis. Region-wise BOLD signal time series were extracted from a whole-brain parcellation scheme (e.g., the Glasser atlas) and whole-brain effective connectivity was inferred using rDCM. The rDCM parameter estimates were then analysed with regard to group-level consistency and test-retest reliability. **(B)** Estimates of the probability density functions (using the nonparametric kernel smoothing of *fitdist.m* implemented in MATLAB) of the connection-wise intra-class correlation coefficient (ICC) for the resting-state and all 7 tasks (i.e., emotional processing, gambling, language, motor, relational processing, social cognition, and working memory) for the Glasser atlas (see Supplementary Figure S1 for the respective results of the Schaefer atlas). Results are shown when considering all connections (*left*), significant connections (*middle*), and the top-1000 connections (*right*). **(C)** Mean (averaged across all paradigms) test-retest reliability for all connections (*top, left*) as well as how often (i.e., in how many paradigms) a connection showed excellent reliability (*bottom, left*). Mean test-retest reliability projected onto the cortical surface (*top, middle*) and the cortical location of all regions that are linked via connections that show excellent reliability in all 8 paradigms (*bottom, middle*). Connectogram showing the connections with excellent reliability in all 8 paradigms (*right*). The connectogram was produced using Circos (publicly available at http://circos.ca/software/).

Focusing only on connections that deviated significantly from zero (*p* < 0.05, false discovery rate (FDR)-corrected for multiple comparisons), we observed a clear increase in reliability (**Fig. 1B**, *middle*). While reliability of the significant connections inferred from resting-state fMRI data was still poor on average (mean ICC = 0.32 [-0.10 0.64]), reliability was considerably higher for task-based fMRI data (e.g., mean ICC = 0.62 [0.10 0.88] for the emotional processing task). The same pattern could be observed for the significant driving inputs although (somewhat less strongly). Finally, when restricting our reliability analysis even further to the top-1000 connections (i.e., the connections with the highest absolute connection strengths), we found the shift towards higher reliability to be even more pronounced (**Fig. 1B**, *right*). Specifically, we found reliability to range on average from fair for the resting state (mean ICC = 0.45 [-0.02 0.76]) to near-excellent for the emotional processing task (mean ICC = 0.74 [0.45 0.89]). A comprehensive list of all results from the test-retest reliability analysis is provided in **Table 2**.

**Table 2.**
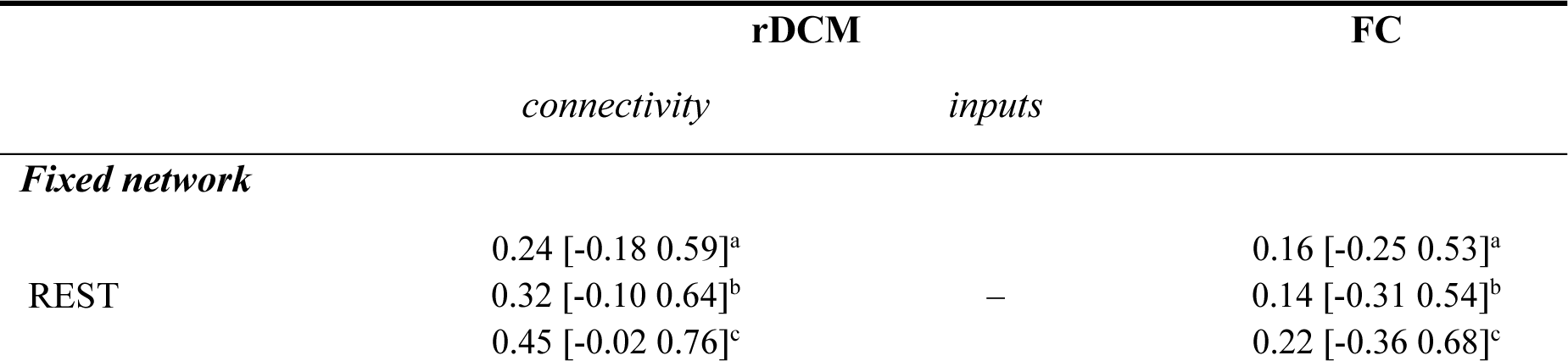
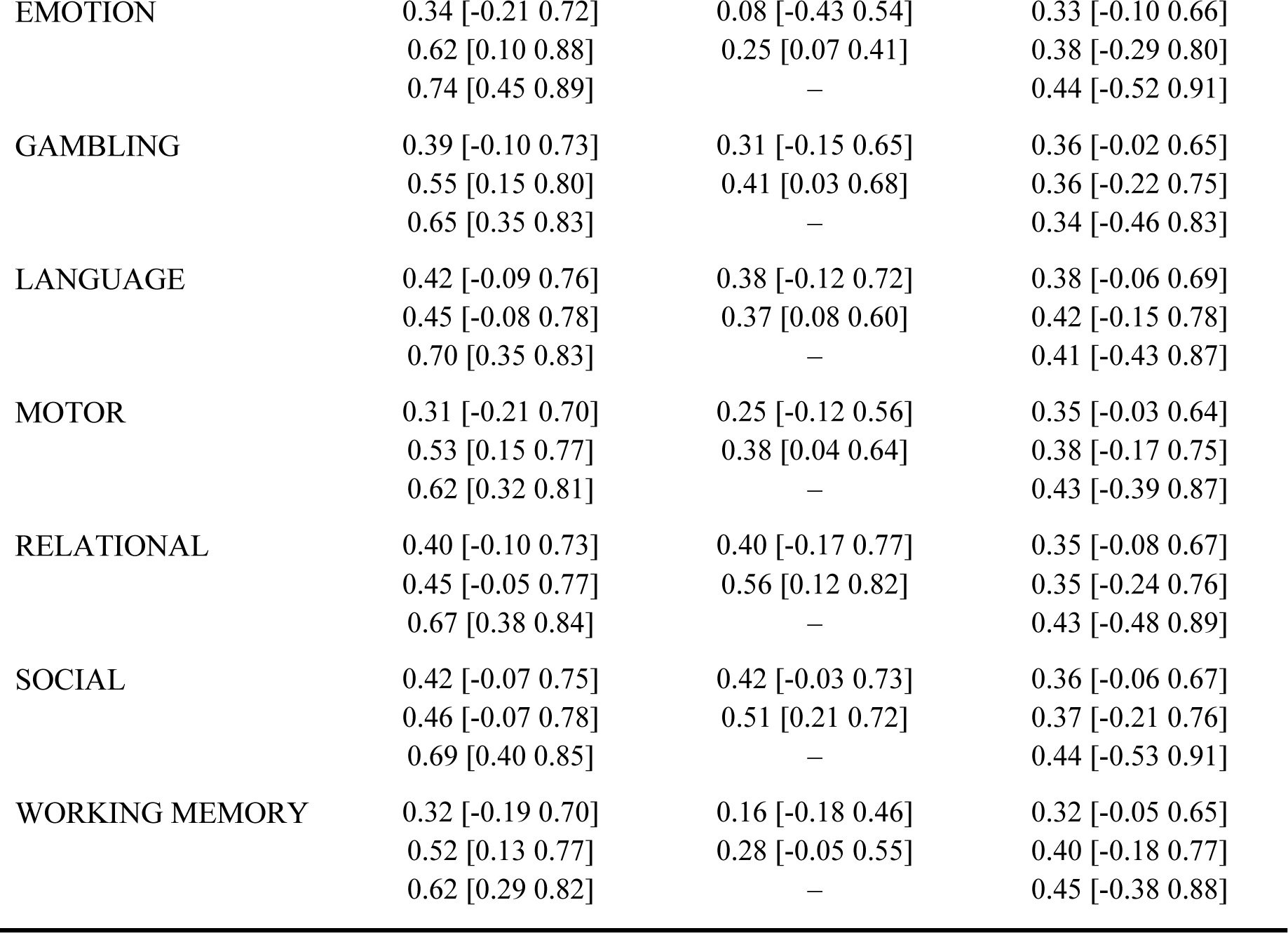
Test-retest reliability of model parameter estimates for regression DCM and functional connectivity. Test-retest reliability of parameter estimates was assessed in terms of the intra-class correlation coefficient (ICC) between estimates of session 1 (“test”) and session 2 (“retest”) for a fixed (full) network architecture (i.e., classical rDCM and Pearson correlation coefficient). Here, we report the mean (averaged across parameters) ICC value and 95% confidence interval (CI). Averaging of the connection-wise ICC values as well as computing the 95% CI was achieved by (i) transforming connection-wise ICC values to z-space using Fisher-z transformation, (ii) computing mean as well as lower and upper bound of the 95% CI in z-space, and finally (iii) back-transforming estimates to r-space. Test-retest reliability is reported for the connectivity and driving input parameter estimates of rDCM (*middle*) as well as for the functional connectivity estimates (*right*). For both methods, results are shown for all HCP paradigms for the Glasser atlas (see Supplementary Table S1 for the respective results of the Schaefer atlas). Furthermore, results are shown for: (^a^) all parameters (*top row*), (^b^) significant parameters (*middle row*), and (^c^) top-1000 parameters (*bottom row*).

|  | rDCM |  | FC |
| --- | --- | --- | --- |
|  | connectivity | inputs |  |
| **Fixed network** |  |  |  |
| REST | 0.24 [-0.18 0.59]<sup>a</sup> |  | 0.16 [-0.25 0.53]<sup>a</sup> |
|  | 0.32 [-0.10 0.64]<sup>b</sup> | — | 0.14 [-0.31 0.54]<sup>b</sup> |
|  | 0.45 [-0.02 0.76]<sup>c</sup> |  | 0.22 [-0.36 0.68]<sup>c</sup> |
| EMOTION | 0.34 [-0.21 0.72] | 0.08 [-0.43 0.54] | 0.33 [-0.10 0.66] |
|  | 0.62 [0.10 0.88] | 0.25 [0.07 0.41] | 0.38 [-0.29 0.80] |
|  | 0.74 [0.45 0.89] | – | 0.44 [-0.52 0.91] |
| GAMBLING | 0.39 [-0.10 0.73] | 0.31 [-0.15 0.65] | 0.36 [-0.02 0.65] |
|  | 0.55 [0.15 0.80] | 0.41 [0.03 0.68] | 0.36 [-0.22 0.75] |
|  | 0.65 [0.35 0.83] | – | 0.34 [-0.46 0.83] |
| LANGUAGE | 0.42 [-0.09 0.76] | 0.38 [-0.12 0.72] | 0.38 [-0.06 0.69] |
|  | 0.45 [-0.08 0.78] | 0.37 [0.08 0.60] | 0.42 [-0.15 0.78] |
|  | 0.70 [0.35 0.83] | – | 0.41 [-0.43 0.87] |
| MOTOR | 0.31 [-0.21 0.70] | 0.25 [-0.12 0.56] | 0.35 [-0.03 0.64] |
|  | 0.53 [0.15 0.77] | 0.38 [0.04 0.64] | 0.38 [-0.17 0.75] |
|  | 0.62 [0.32 0.81] | – | 0.43 [-0.39 0.87] |
| RELATIONAL | 0.40 [-0.10 0.73] | 0.40 [-0.17 0.77] | 0.35 [-0.08 0.67] |
|  | 0.45 [-0.05 0.77] | 0.56 [0.12 0.82] | 0.35 [-0.24 0.76] |
|  | 0.67 [0.38 0.84] | – | 0.43 [-0.48 0.89] |
| SOCIAL | 0.42 [-0.07 0.75] | 0.42 [-0.03 0.73] | 0.36 [-0.06 0.67] |
|  | 0.46 [-0.07 0.78] | 0.51 [0.21 0.72] | 0.37 [-0.21 0.76] |
|  | 0.69 [0.40 0.85] | – | 0.44 [-0.53 0.91] |
| WORKING MEMORY | 0.32 [-0.19 0.70] | 0.16 [-0.18 0.46] | 0.32 [-0.05 0.65] |
|  | 0.52 [0.13 0.77] | 0.28 [-0.05 0.55] | 0.40 [-0.18 0.77] |
|  | 0.62 [0.29 0.82] | – | 0.45 [-0.38 0.88] |

In a post-hoc analysis, we inspected which connections were most reliable across the different HCP paradigms. The mean ICC values (averaged across all eight paradigms) revealed a notable pattern of connections that were consistently reliable across paradigms (**Fig. 1C**, *left*). In particular, when inspecting connections that showed excellent reliability (i.e., ICC > 0.75) in all eight paradigms, we found these connections to primarily link regions such as areas a9-46v, a47r, p47r, and p10p near the frontal pole, AVI and FOP5 in the anterior insula and the frontal operculum, respectively, as well as TE1m and TE2a on the lateral surface of the temporal lobe (**Fig. 1C**, *bottom left*). These regions map well onto components of the multiple-demands network, which is characterized by showing consistent activation for a number of different cognitive tasks (Assem et al., 2020; Fedorenko et al., 2013).

These results illustrate that stronger connections (both inhibitory and excitatory) inferred by rDCM are more reliable across sessions and, in fact, often achieve good to excellent test-retest reliability (i.e., ICC > 0.6). This is confirmed when directly testing the correlation between the absolute mean (i.e., averaged across all participants) parameter strength and the ICC value of the parameter estimate, both for connectivity parameters (for all paradigms: *r* ≥ 0.26, all *p* < 0.001) and driving input parameters – although this was more variable for the latter (range: *r* = -0.04, *p* = 0.29 to *r* = 0.40, *p* < 0.001).

#### 3.2.2 Regression DCM with sparsity constraints

In a second step, we assessed the test-retest reliability of connectivity estimates obtained using rDCM with embedded sparsity constraints. Overall, the test-retest reliability of parameter estimates from sparse rDCM was lower than for rDCM with fixed network architecture.

When considering all connections, test-retest reliability was on average poor for all paradigms (**Fig. 2A**, *left*). More specifically, for the connectivity parameters, test-retest reliability ranged from mean ICC = 0.02 [-0.28 0.33] for the resting state to mean ICC = 0.34 [-0.09 0.66] for the motor task when considering all connections. Similarly, for the driving input parameters, test-retest reliability ranged from poor for the motor task (mean ICC = 0.11 [-0.24 0.44]) to fair for the relational processing task (mean ICC = 0.40 [-0.16 0.76]).

**Figure 2:**
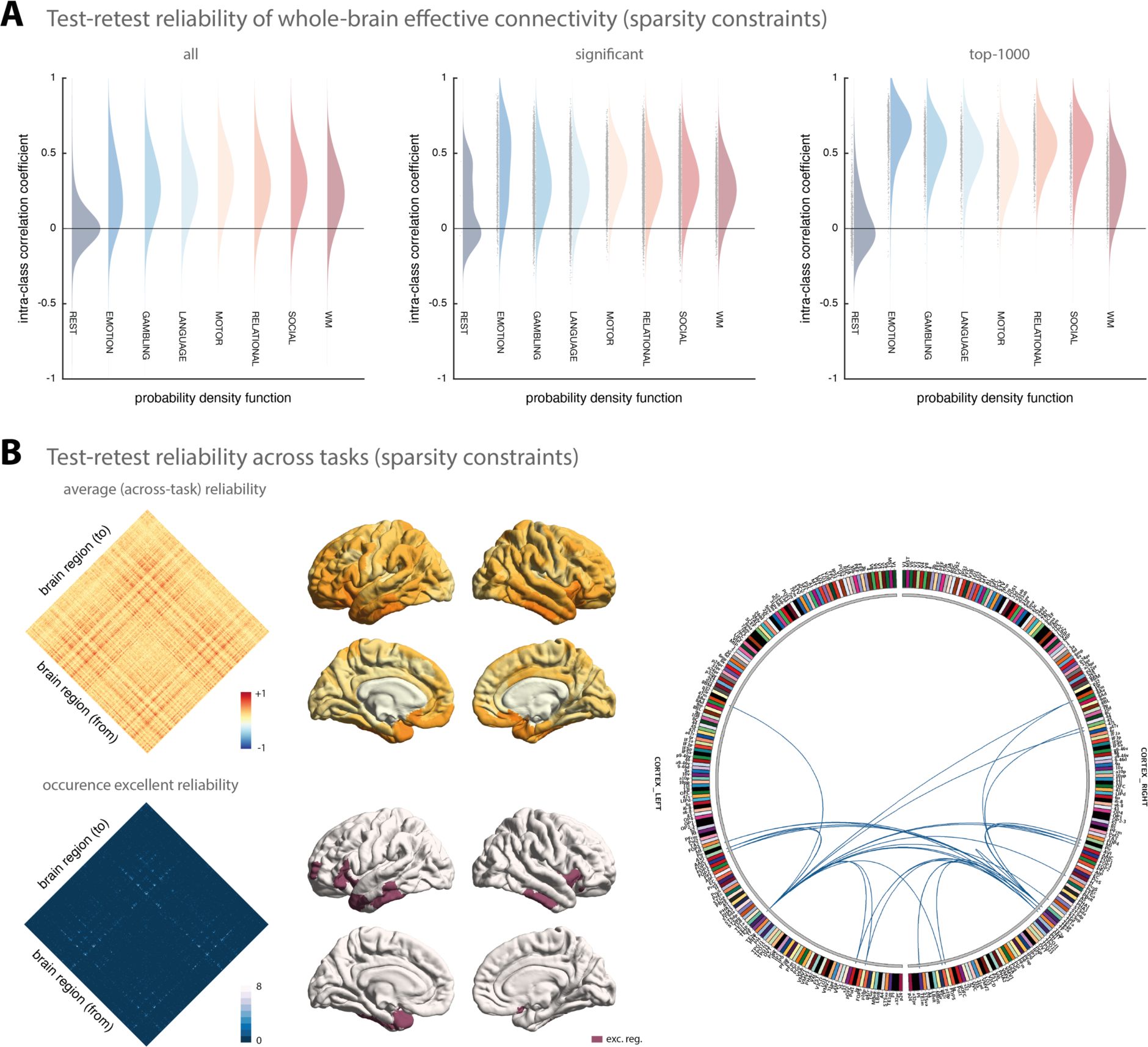
Test-retest reliability of regression DCM with sparsity constraints. **(A)** Estimates of the probability density functions (using the nonparametric kernel smoothing of *fitdist.m* implemented in MATLAB) of the connection-wise intra-class correlation coefficient (ICC) for the resting-state and all 7 tasks (i.e., emotional processing, gambling, language, motor, relational processing, social cognition, and working memory) for the Glasser atlas (see Supplementary Figure S2 for the respective results of the Schaefer atlas). Results are shown when considering all connections (*left*), significant connections (*middle*), and the top-1000 connections (*right*). **(B)** Mean (averaged across all paradigms) test-retest reliability for all connections (*top, left*) as well as how often (i.e., in how many paradigms) a connection showed excellent reliability (*bottom, left*). Mean test-retest reliability projected onto the cortical surface (*top, middle*) and the cortical location of all regions that are linked via connections that show excellent reliability in at least 6 paradigms (*bottom, middle*). Connectogram showing the connections with excellent reliability in at least 6 paradigms (*right*). The connectogram was produced using Circos (publicly available at http://circos.ca/software/).

In a next step, we again tested whether stronger connections were more reliable. Focusing only on connections that deviated significantly from zero (*p* < 0.05, FDR-corrected), we again observed a shift towards higher reliability (**Fig. 2A**, *middle*), although less pronounced as for rDCM with fixed network architecture. For sparse rDCM, reliability of the significant connectivity parameters ranged on average from poor for the resting state (mean ICC = 0.16 [-0.32 0.57]) to fair for the emotional processing task (mean ICC = 0.44 [-0.18 0.81]). The same pattern could be observed for the significant driving input estimates. Finally, when restricting our reliability analysis even further to the top-1000 connections, we found the shift towards higher reliability to be even more pronounced – with one exception: the resting state (**Fig. 2A**, *right*). Specifically, even for the top 1000 connections, we found poor reliability for the resting state (mean ICC = 0.05 [-0.29 0.38]), whereas for all task-based fMRI datasets, test-retest reliability was considerably increased when considering only the top-1000 connections (e.g., mean ICC = 0.66 [0.25 0.86] for the emotional processing task). A comprehensive list of all results from the test-retest reliability analysis for sparse rDCM is provided in **Table 3**.

**Table 3.** Test-retest reliability of model parameter estimates for regression DCM and functional connectivity (sparsity constraints). Test-retest reliability of parameter estimates was assessed in terms of the intra-class correlation coefficient (ICC) between estimates of session 1 (“test”) and session 2 (“retest”) for sparsity constraints (i.e., rDCM with sparsity constraints and L1-regularized partial correlations). Here, we report the mean (averaged across parameters) ICC value and 95% confidence interval (CI). Averaging of the connection-wise ICC values as well as computing the 95% CI was done in z-space (see caption of Table 2 for details). Test-retest reliability is reported for the connectivity and driving input estimates of rDCM (*middle*) as well as for the functional connectivity estimates (*right*). For both methods, results are shown for all HCP paradigms for the Glasser atlas (see Supplementary Table S2 for the respective results of the Schaefer atlas). Furthermore, results are shown for: (^a^) all parameters (*top row*), (^b^) significant parameters (*middle row*), and (^c^) top-1000 parameters (*bottom row*).

|  | rDCM |  | FC |
| --- | --- | --- | --- |
|  | *connectivity* | *inputs* |  |
| ***Fixed network*** |  |  |  |
| REST | 0.02 [-0.28 0.33]<sup>a</sup> |  | 0.14 [-0.38 0.60]<sup>a</sup> |
|  | 0.16 [-0.32 0.57]<sup>b</sup> | – | 0.50 [0.02 0.79]<sup>b</sup> |
|  | 0.05 [-0.29 0.38]<sup>c</sup> |  | 0.55 [0.11 0.81]<sup>c</sup> |
| EMOTION | 0.25 [-0.24 0.64] | 0.20 [-0.26 0.54] | 0.08 [-0.44 0.55] |
|  | 0.44 [-0.18 0.81] | 0.73 [0.73 0.73] | 0.30 [-0.07 0.59] |
|  | 0.66 [0.25 0.86] | – | 0.31 [-0.10 0.63] |
| GAMBLING | 0.29 [-0.15 0.63] | 0.26 [-0.20 0.63] | 0.08 [-0.42 0.54] |
|  | 0.31 [-0.12 0.65] | 0.42 [0.01 0.70] | 0.32 [-0.06 0.61] |
|  | 0.56 [0.20 0.79] | – | 0.32 [-0.04 0.61] |
| LANGUAGE | 0.27 [-0.14 0.61] | 0.38 [-0.12 0.73] | 0.08 [-0.43 0.56] |
|  | 0.29 [-0.10 0.61] | 0.33 [-0.02 0.61] | 0.36 [-0.06 0.66] |
|  | 0.51 [0.14 0.76] | – | 0.37 [-0.05 0.68] |
| MOTOR | 0.34 [-0.09 0.66] | 0.11 [-0.24 0.44] | 0.07 [-0.43 0.54] |
|  | 0.38 [0.02 0.65] | – | 0.33 [-0.03 0.62] |
|  | 0.43 [0.06 0.69] | – | 0.35 [-0.06 0.66] |
| RELATIONAL | 0.30 [-0.13 0.64] | 0.40 [-0.16 0.76] | 0.07 [-0.41 0.52] |
|  | 0.33 [-0.07 0.64] | 0.54 [0.08 0.81] | 0.34 [-0.02 0.62] |
|  | 0.55 [0.21 0.78] | – | 0.33 [-0.07 0.64] |
| SOCIAL | 0.31 [-0.11 0.64] | 0.36 [-0.09 0.69] | 0.09 [-0.46 0.59] |
|  | 0.33 [-0.07 0.63] | 0.46 [0.13 0.70] | 0.40 [0.03 0.67] |
|  | 0.56 [0.19 0.79] | – | 0.40 [0.00 0.69] |
| WORKING MEMORY | 0.24 [-0.15 0.57] | 0.16 [-0.20 0.48] | 0.08 [-0.38 0.51] |
|  | 0.26 [-0.09 0.56] | 0.27 [0.14 0.39] | 0.35 [0.00 0.63] |
|  | 0.32 [-0.06 0.62] | – | 0.37 [0.00 0.64] |

In a post-hoc analysis, we again inspected which connections were most reliable across the different HCP paradigms. Inspecting the mean (averaged across all paradigms) ICC values revealed a similar pattern for sparse rDCM as observed above for classical rDCM – although with somewhat lower mean ICC values (**Fig. 2B**, *left*). For example, no connections were found that showed excellent reliability in all eight paradigms. However, when inspecting those connections that showed excellent reliability in at least 6 of the 8 paradigms, we observed a pattern that was highly consistent to the one obtained using rDCM with fixed network architecture (see above). Specifically, these connections again primarily linked regions that had previously been identified with the multiple-demands network, such as areas p10p near the frontal pole, AVI and FOP5 in the anterior insula and the frontal operculum, respectively, as well as TE1m and TE2a on the lateral surface of the temporal lobe (**Fig. 2B**, *bottom left*).

Again, these results illustrate that stronger parameters (both inhibitory and excitatory) are more reliable than weaker parameters. This observation was confirmed when explicitly testing the correlation between the mean (i.e., averaged across all participants) absolute parameter strength and the ICC values of the parameter estimate, both for connectivity strengths (resting state: *r* = 0.01, *p* < 0.001; for all task paradigms: *r* ≥ 0.18, *p* < 0.001) and driving inputs, although this was again more variable for the latter (range: *r* = 0.09; *p* = 0.01 to *r* = 0.39, *p* < 0.001).

In summary, our results indicate that, for the present datasets, connectivity estimates obtained using sparse rDCM were less reliable than those obtained using rDCM with fixed network architecture (see Discussion for potential explanations). For resting-state data, test-retest reliability of sparse rDCM was poor – even when focusing on strong connections. Conversely, for the driving input estimates, test-retest reliability was comparable across the two rDCM variants.

#### 3.2.3 Comparison to functional connectivity

For comparison with rDCM, we investigated the test-retest reliability of functional connectivity estimates obtained using Pearson correlations and L1-regularized partial correlations.

First, we compared results from rDCM with fixed network architecture to Pearson correlations (**Fig. 3A)**. We found that the two methods showed similar test-retest reliability when considering all model parameters (**Fig. 3A**, *left*). Specifically, test-retest reliability of Pearson correlations ranged from mean ICC = 0.16 [-0.25 0.53] for the resting state to mean ICC = 0.38 [-0.06 0.69] for the language task. Interestingly, when focusing on stronger connections, Pearson correlations did not show the same improvement previously observed for rDCM; instead, test-retest reliability remained mostly poor (or fair at best). More specifically, when focusing only on significant parameter estimates, reliability ranged from mean ICC = 0.14 [-0.31 0.54] for the resting state to mean ICC = 0.42 [-0.15 0.78] for the language task (**Fig. 3A**, *middle*). Similarly, when restricting the analysis to the top-1000 connections, reliability ranged from mean ICC = 0.22 [-0.36 0.68] for the resting state to mean ICC = 0.45 [-0.38 0.88] for the working memory task (**Fig. 3A**, *right*). A comprehensive list of all results from the test-retest reliability analysis is provided in **Table 2** (*right column*).

**Figure 3:**
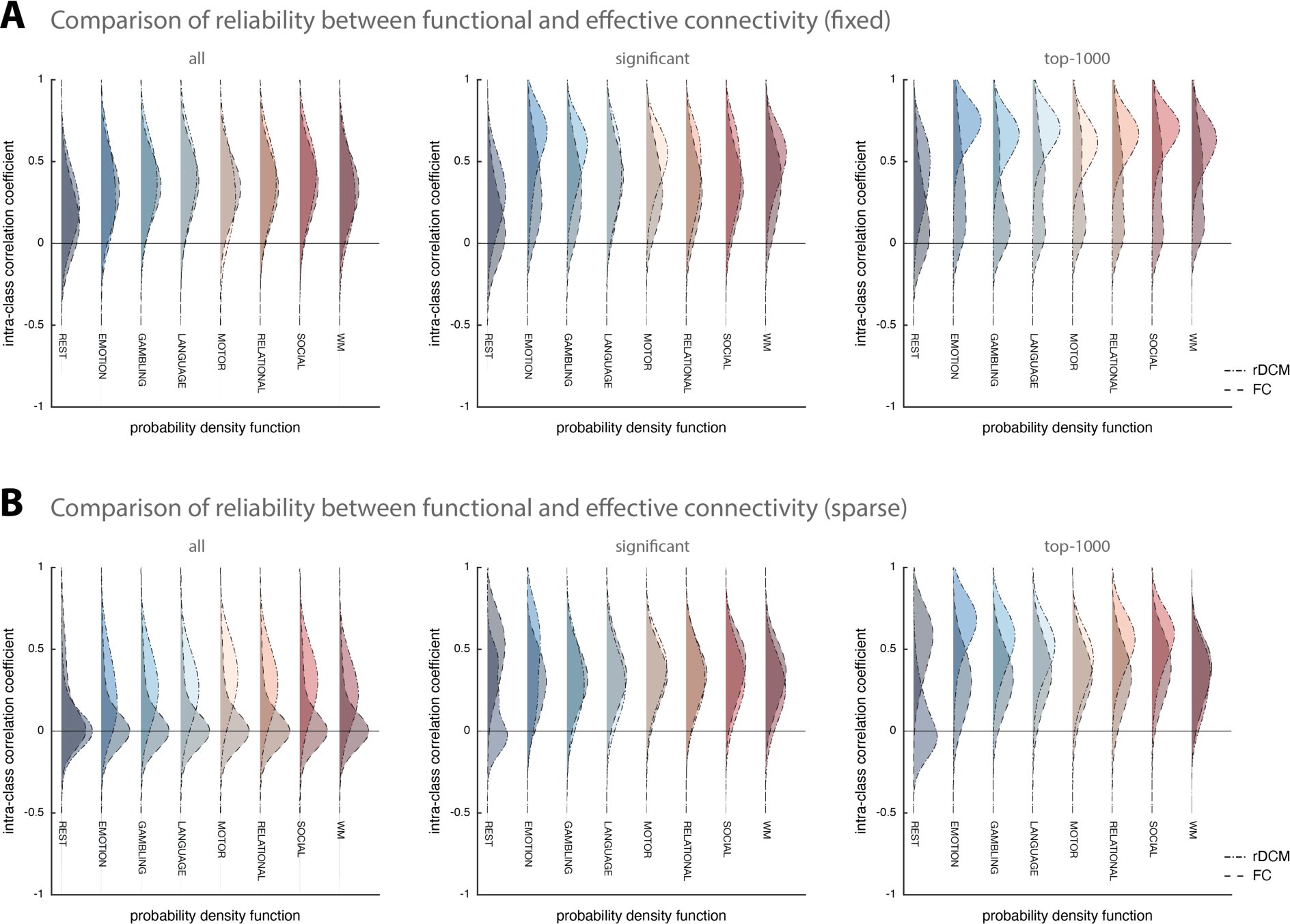
Comparison of test-retest reliability between regression DCM and functional connectivity. **(A)** Estimates of the probability density functions (using the nonparametric kernel smoothing of *fitdist.m* implemented in MATLAB) of the connection-wise intra-class correlation coefficient (ICC) for the resting-state and all 7 tasks (i.e., emotional processing, gambling, language, motor, relational processing, social cognition, and working memory) for the Glasser atlas (see Supplementary Figure S3 for the respective results of the Schaefer atlas) for fixed (full) connectivity methods (i.e., classical rDCM and Pearson correlation coefficient), and **(B)** sparse connectivity methods (i.e., rDCM with sparsity constraints and L1-regularized partial correlations). Probability density functions representing rDCM results are shown with dot-dashed lines and lighter colors, whereas probability density functions representing functional connectivity results are shown with dashed lines and darker colors. For each connectivity variant, results are shown when considering all connections (*left*), significant connections (*middle*), and the top-1000 connections (*right*).

Second, we compared sparse rDCM to L1-regularized partial correlations (**Fig. 3B)**. Interestingly, we found test-retest reliability of L1-regularized partial correlations to be on average close-to-zero for all paradigms when considering all connectivity parameters (**Fig. 3B**, *left*). Specifically, test-retest reliability ranged from mean ICC = 0.07 [-0.41 0.52] for the relational processing task to mean ICC = 0.14 [-0.38 0.60] for the resting state. While this improved when focusing on stronger connections, test-retest reliability remained relatively low for L1-regularized partial correlations and, in most cases, worse than for sparse rDCM. More specifically, when focusing only on significant parameters, reliability ranged from mean ICC = 0.30 [-0.07 0.59] for the emotional processing task to mean ICC = 0.50 [0.02 0.79] for the resting state (**Fig. 3B**, *middle*). Similarly, when restricting the analysis to the top-1000 connections, reliability ranged from mean ICC = 0.31 [-0.10 0.63] for the emotional processing task to mean ICC = 0.55 [0.11 0.81] for the resting state (**Fig. 3B**, *right*). A comprehensive list of all results from the test-retest reliability analysis is provided in **Table 3** (*right column*).

### 3.3 Similarity analysis: inter-session consistency of whole-brain effective connectivity patterns

#### Regression DCM with fixed (fully connected) network architecture

In a final analysis, we shifted the focus from reliability of separate connections to the consistency of the whole-brain effective connectivity profile across time. To this end, we asked whether the effective connectivity profile of an individual person obtained in one session could be used to identify this individual from the set of all effective connectivity profiles obtained in another session. This analysis follows previous work demonstrating that functional connectivity profiles derived from fMRI data enables the identification of individual subjects (Finn et al., 2015).

First, we assessed identification accuracies for the whole-brain effective connectivity patterns inferred using rDCM with fixed network architecture (chance level: 1/N_sub_ * 100%, ranging from 2.4% to 2.3%, depending on the number of subjects available in each task). Overall, entire effective connectivity profiles were highly consistent across the two sessions and enabled identification of individual participants with high accuracies. More specifically, when predicting identity in session 2 from session 1 (S_1_ → S_2_), identification accuracies ranged from 80.5% (33/41) for the emotional processing task to 100% (44/44) for the social cognition task. Similarly, when predicting identity in session 1 from session 2 (S_2_ → S_1_), identification accuracies ranged from 78.0% (32/41) for the emotional processing task to 100% (44/44) for the social cognition task. Results were almost identical for the Schaefer parcellation. All of the identification accuracies were statistically significant at *p* < 0.05 (Bonferroni-corrected for multiple comparisons), as assessed using permutation testing (see Methods). A comprehensive list of all identification accuracies is provided in **Table 4** (*middle column, top*).

**Table 4.**
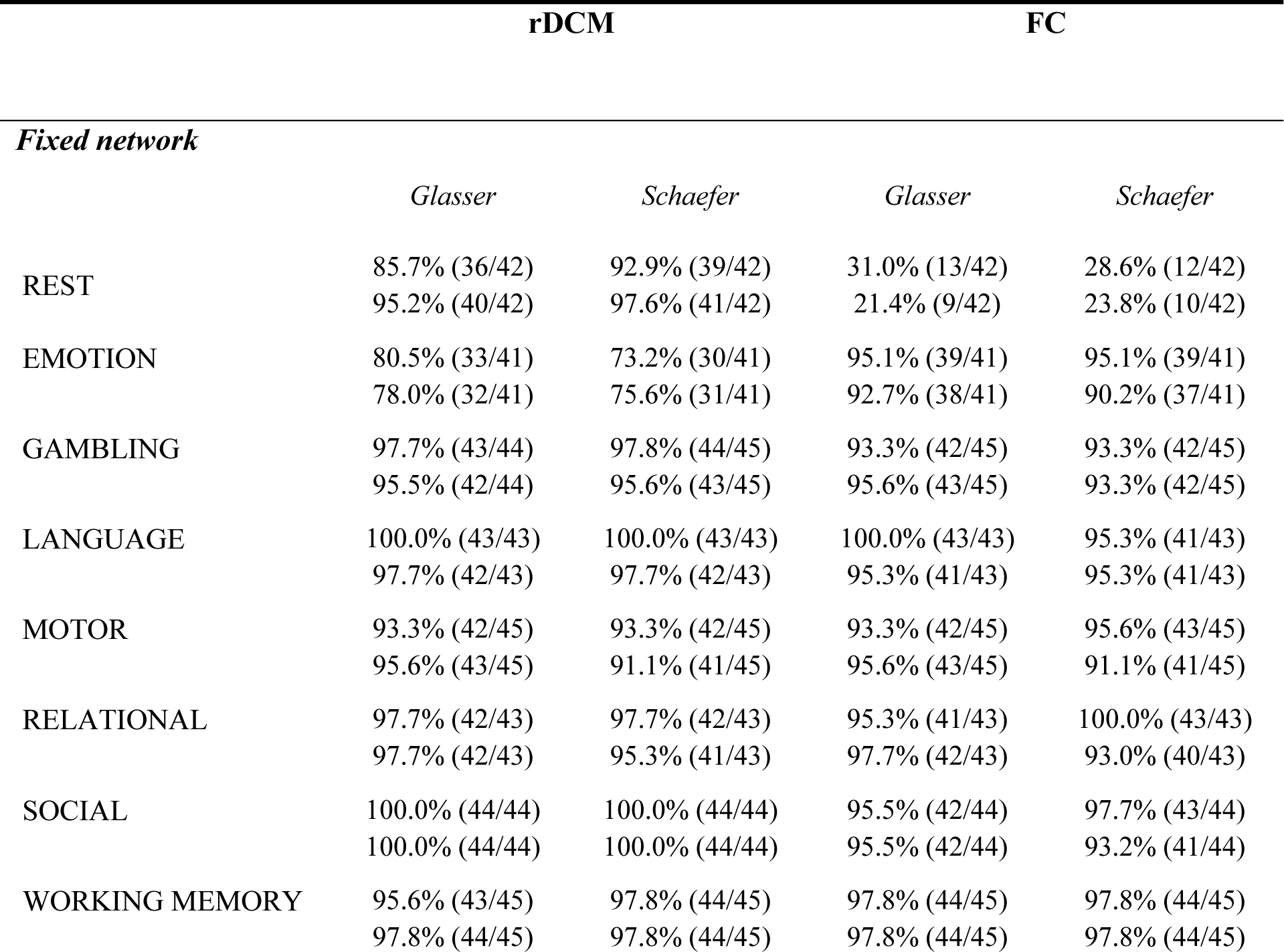
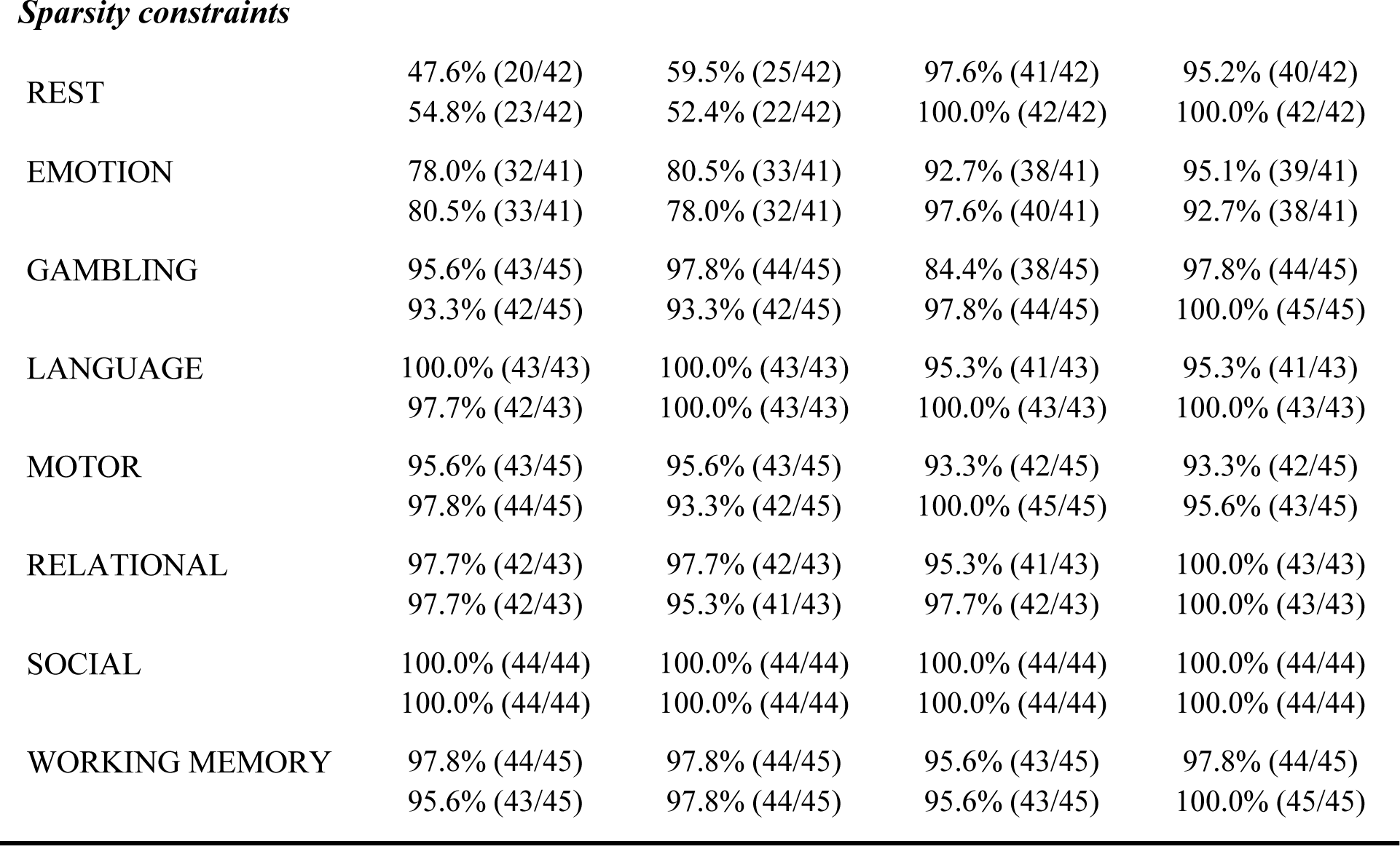
Across-session consistency of connectivity profiles for regression DCM and functional connectivity. Consistency of the entire connectivity profile across the two sessions. Identification accuracies are reported for predicting identity in session 2 from session 1 (*top*) and vice versa (*bottom*). Results are reported for rDCM (*middle*) and for functional connectivity estimates (*right*). For both methods, results are shown for all HCP paradigms as well as for the two whole-brain parcellation schemes (i.e., Glasser, Schaefer). Furthermore, results are reported for two different “modes” of estimation (see main text for details): (i) fixed network architecture (i.e., classical rDCM and Pearson correlation coefficient), and (ii) sparsity constraints (i.e., rDCM with sparsity constraints and L1-regularized partial correlations).

|  | rDCM |  | FC |  |
| --- | --- | --- | --- | --- |
| *Fixed network* |  |  |  |  |
|  | *Glasser* | *Schaefer* | *Glasser* | *Schaefer* |
| REST | 85.7% (36/42) | 92.9% (39/42) | 31.0% (13/42) | 28.6% (12/42) |
|  | 95.2% (40/42) | 97.6% (41/42) | 21.4% (9/42) | 23.8% (10/42) |
| EMOTION | 80.5% (33/41) | 73.2% (30/41) | 95.1% (39/41) | 95.1% (39/41) |
|  | 78.0% (32/41) | 75.6% (31/41) | 92.7% (38/41) | 90.2% (37/41) |
| GAMBLING | 97.7% (43/44) | 97.8% (44/45) | 93.3% (42/45) | 93.3% (42/45) |
|  | 95.5% (42/44) | 95.6% (43/45) | 95.6% (43/45) | 93.3% (42/45) |
| LANGUAGE | 100.0% (43/43) | 100.0% (43/43) | 100.0% (43/43) | 95.3% (41/43) |
|  | 97.7% (42/43) | 97.7% (42/43) | 95.3% (41/43) | 95.3% (41/43) |
| MOTOR | 93.3% (42/45) | 93.3% (42/45) | 93.3% (42/45) | 95.6% (43/45) |
|  | 95.6% (43/45) | 91.1% (41/45) | 95.6% (43/45) | 91.1% (41/45) |
| RELATIONAL | 97.7% (42/43) | 97.7% (42/43) | 95.3% (41/43) | 100.0% (43/43) |
|  | 97.7% (42/43) | 95.3% (41/43) | 97.7% (42/43) | 93.0% (40/43) |
| SOCIAL | 100.0% (44/44) | 100.0% (44/44) | 95.5% (42/44) | 97.7% (43/44) |
|  | 100.0% (44/44) | 100.0% (44/44) | 95.5% (42/44) | 93.2% (41/44) |
| WORKING MEMORY | 95.6% (43/45) | 97.8% (44/45) | 97.8% (44/45) | 97.8% (44/45) |
|  | 97.8% (44/45) | 97.8% (44/45) | 97.8% (44/45) | 97.8% (44/45) |
| *Sparsity constraints* |  |  |  |  |
| REST | 47.6% (20/42) | 59.5% (25/42) | 97.6% (41/42) | 95.2% (40/42) |
|  | 54.8% (23/42) | 52.4% (22/42) | 100.0% (42/42) | 100.0% (42/42) |
| EMOTION | 78.0% (32/41) | 80.5% (33/41) | 92.7% (38/41) | 95.1% (39/41) |
|  | 80.5% (33/41) | 78.0% (32/41) | 97.6% (40/41) | 92.7% (38/41) |
| GAMBLING | 95.6% (43/45) | 97.8% (44/45) | 84.4% (38/45) | 97.8% (44/45) |
|  | 93.3% (42/45) | 93.3% (42/45) | 97.8% (44/45) | 100.0% (45/45) |
| LANGUAGE | 100.0% (43/43) | 100.0% (43/43) | 95.3% (41/43) | 95.3% (41/43) |
|  | 97.7% (42/43) | 100.0% (43/43) | 100.0% (43/43) | 100.0% (43/43) |
| MOTOR | 95.6% (43/45) | 95.6% (43/45) | 93.3% (42/45) | 93.3% (42/45) |
|  | 97.8% (44/45) | 93.3% (42/45) | 100.0% (45/45) | 95.6% (43/45) |
| RELATIONAL | 97.7% (42/43) | 97.7% (42/43) | 95.3% (41/43) | 100.0% (43/43) |
|  | 97.7% (42/43) | 95.3% (41/43) | 97.7% (42/43) | 100.0% (43/43) |
| SOCIAL | 100.0% (44/44) | 100.0% (44/44) | 100.0% (44/44) | 100.0% (44/44) |
|  | 100.0% (44/44) | 100.0% (44/44) | 100.0% (44/44) | 100.0% (44/44) |
| WORKING MEMORY | 97.8% (44/45) | 97.8% (44/45) | 95.6% (43/45) | 97.8% (44/45) |
|  | 95.6% (43/45) | 97.8% (44/45) | 95.6% (43/45) | 100.0% (45/45) |

#### Regression DCM with sparsity constraints

Second, identification accuracies were assessed for sparse rDCM. Again, the sparse whole-brain effective connectivity profiles were highly consistent across the two sessions and allowed identification of individual participants with high accuracies, with the notable exception of the resting state. More specifically, for the resting state, identification accuracies were around 50% (i.e., 47.6% when predicting S_1_ → S_2_, and 54.8% when predicting S_2_ → S_1_); please see **Table 4** for details. For task-based data, identification accuracies were considerably higher. Specifically, when predicting S_1_ →S_2_, identification accuracies ranged from 78.0% (32/41) for the emotional processing task to 100% (44/44) for the social cognition task. Similarly, when predicting S_2_ → S_1_, identification accuracies ranged from 80.5% (33/41) for the emotional processing task to 100% (44/44) for the social cognition task. Again, all identification accuracies – even for the resting state – were statistically significant at *p* < 0.05 (Bonferroni-corrected), as assessed using permutation testing.

#### Comparison to functional connectivity

Finally, we compared identification accuracies between rDCM and functional connectivity estimates obtained using Pearson correlation and L1-regularized partial correlations. Overall, we found that functional connectivity profiles also enabled identification of individual participants with high accuracies (**Table 4**). There was one notable exception: connectivity during the resting state, as characterized by Pearson correlation coefficients. More specifically, for this setting, identification accuracies were 31.0% (13/42) when predicting S_1_ → S_2_, and 21.4% (9/42) when predicting S_2_ → S_1_. This is in contrast to previous reports by Finn et al. (2015); for potential explanations of these inconsistencies, please see the Discussion. For all other settings, identification accuracies of functional connectivity profiles were high and even surpassed those reported for rDCM in some cases, particularly when using sparsity constraints. Again, all identification accuracies – even for the resting state in combination with Pearson’s correlations – were statistically significant at *p* < 0.05 (Bonferroni-corrected), as assessed using permutation testing.

## 4 Discussion

In this paper, we assessed the test-retest reliability and group-level consistency of connection strengths inferred from fMRI data using rDCM (Frässle et al., 2021b; 2018a; 2017). First, using two different whole-brain parcellations, we demonstrated that rDCM provides highly consistent parameter estimates at the group level across two sessions of the HCP dataset (Van Essen et al., 2013), regardless of the exact paradigm. Second, we found, on average, relatively low test-retest reliability when considering all connections. However, stronger connections were more reliable, with many strong connections displaying good to excellent test-retest reliability (ICC ≥ 0.6); see **Table 2**. When comparing this to the test-retest reliability of measures of functional connectivity, rDCM performed favorably – in particular, when focusing on strong connections (see **Fig. 3**). While these observations hold for both variants of rDCM, we found test-retest reliability to be considerably higher for rDCM with fixed network architecture as compared to rDCM with sparsity constraints.

The increase in reliability with higher connection strengths is worth emphasizing. For example, when restricting the analysis to the top-1000 connections, we found for all task-based datasets on average good test-retest reliability (see **Table 2**). This suggests that those connections representing meaningful effects can be reliably inferred using rDCM. These observations are consistent with previous analyses of test-retest reliability in the context of classical DCM for fMRI. For instance, Frässle et al. (2016) assessed test-retest reliability of effective connectivity in small (six-region) networks of the core face perception system. While finding fair-to-good reliability of parameter estimates on average, they observed a similar trend of increased reliability for larger parameter estimates. Our results are also in line with other reports on the test-retest reliability of classical DCM (Frässle et al., 2015; Rowe et al., 2010; Schuyler et al., 2010) and spectral DCM (Almgren et al., 2018) – all conducted in the context of much smaller networks than the ones considered here. Furthermore, the observed increase in test-retest reliability with connection strength is not exclusive to DCMs. For instance, a similar increase of test-retest reliability with effect size has also been observed in conventional fMRI analyses (Caceres et al., 2009).

Interestingly, this pattern of increased test-retest reliability for stronger connections was less pronounced for functional connectivity estimates (**Fig. 3**). Test-retest reliability estimates based on Pearson correlations and L1-regularized partial correlations also an increase of ICC values for greater connection strength, in line with previous studies of functional connectivity (for review, see Noble et al., 2019). However, this increase was only moderate and the average test-retest reliability remained poor to fair, even for the strong connections.

With regard to the test-retest reliability of rDCM, two further observations are worth highlighting. First, we found connectivity estimates from task-based fMRI data to be consistently more reliable than those from resting-state fMRI data. This is remarkable given that resting-state measurements were considerably longer than task measurements, with longer scanning sessions typically being associated with increased reliability (Birn et al., 2013; Noble et al., 2017). More specifically, while (per session) approximately 1 hour of resting-state fMRI data were collected (combined across the phase-encoding directions), task-based fMRI data comprised just a couple of minutes. Despite these very short scanning sessions, task-based fMRI exhibited superior reliability compared to resting state data. These observations are in line with previous reports demonstrating higher test-retest reliability for functional connectivity patterns derived from task-based as compared to resting-state fMRI data (Noble et al., 2019; Wang et al., 2017). Furthermore, our results are also consistent with findings suggesting that connectivity patterns derived from task-based fMRI are more predictive of individual traits (Greene et al., 2018; 2020). This indicates that – despite its patient-friendly nature – the resting state may not be ideally suited for clinical settings since test-retest reliability is considerably lower than for task-based fMRI – even at much longer scanning times.

Second, we found connectivity estimates by rDCM to be more reliable when assuming a fixed (fully connected) network architecture as compared to relying on embedded sparsity constraints. This was surprising given that sparsity constraints prevent overfitting and should thus increase generalizability of parameter estimates. Having said this, previous simulations have shown that rDCM with sparsity constraints is even more demanding in terms of data quality than rDCM with fixed network architecture (Frässle et al., 2018a). More specifically, we have demonstrated that for low SNR or long TR settings, rDCM with sparsity constraints tends to yield overly sparse connectivity matrices that result from a propensity to pruning existing connections (Frässle et al., 2018a). This may be an explanation for the diminished test-retest reliability observed in the current study in the sense that weak connections may sometimes be pruned and sometimes not.

Finally, moving from assessments of individual connections to whole-brain patterns, we demonstrate that the entire connectivity profile (i.e., the whole-brain “connectivity fingerprint”) of individuals is highly consistent across the two sessions – both for effective (rDCM) and functional connectivity measures. We show that, in many cases, it is possible to identify an individual amongst all participants with close-to-perfect accuracy based on the inferred connectivity pattern. This is consistent with a previous study demonstrating the identifiability of single subjects from functional connectivity measures (Finn et al., 2015), as well as similar reports (Cole et al., 2014; Horien et al., 2019; Noble et al., 2017; Pannunzi et al., 2017; Smith et al., 2009). Interestingly, we found that one particular combination (i.e., resting state and Pearson correlations) yielded relatively low (yet still significant) identification accuracies. This is in contrast to the previous report by Finn et al. (2015). These differences may be due to a number of reasons, including differences in (i) the exact dataset, (ii) preprocessing strategy, or (iii) whole-brain parcellation scheme. Despite this discrepancy, our results support the idea that individual participants may possess unique whole-brain connectivity profile for a given cognitive context. This underscores the exciting opportunities of whole-brain connectivity assessments for studying individual variability of brain networks and how this relates to cognitive phenotypes in health and disease.

Importantly, we here show that all three metrics considered – group-level consistency and test-retest reliability of individual connections as well as whole-brain connectivity profiles – are almost identical for two state-of-the-art parcellation schemes, i.e., the Glasser parcellation (HCP MMP 1.0; Glasser et al., 2016a) and the Schaefer 400 nodes parcellation (Schaefer et al., 2018). This is important because inference on the organizational principles of the brain has been shown to depend on the exact parcellation scheme utilized for defining the nodes of the network (Fornito et al., 2016; 2010). Consequently, it is critical to verify that any conclusions drawn from connectivity estimates are not dependent on this particular choice. Here, we demonstrate that the reliability and consistency of whole-brain effective connectivity estimates obtained using rDCM (as well as those for functional connectivity measures) generalize across the two parcellation schemes.

Our findings have important implications for the fields of human connectomics and network neuroscience in general, as well as for the clinically-oriented disciplines of Computational Psychiatry and Computational Neurology in particular. Especially for the latter two, test-retest reliability of a computational model is important for its clinical utility, particularly when longitudinal measurements are required (e.g., monitoring of treatment response). Here, we showed that rDCM provides good test-retest reliability when focusing on strong connections and enables identification of individual participants with high accuracy based on the entire connectivity profile. Importantly, rDCM shows high reliability even for very short scanning sessions of 3-4 minutes when working with task-based fMRI data. This is important for potential clinical applications.

In summary, our systematic analyses indicate that, in many constellations, rDCM exhibits good properties with regard to group-level consistency and test-retest reliability of connections, as well as the inter-session consistency of whole-brain connectivity patterns. This complements previous methodological assessments of face and construct validity of rDCM (Frässle et al., 2021b; 2018a; 2017; 2021c) and underscores its potential for clinical applications. Its ability to obtain reliable estimates of directed whole-brain connectivity may enable the construction of computational assays for identifying pathophysiological mechanisms and for predictions about individual treatment responses or clinical trajectories (Frässle et al., 2020) – a possibility that we will examine in future studies.

## 5 Code and Data Availability

A Matlab implementation of the regression dynamic causal modeling (rDCM) approach is available as open-source code in the **T**ranslational **A**lgorithms for **P**sychiatry-**A**dvancing **S**cience (TAPAS) software package (www.translationalneuromodeling.org/tapas). Furthermore, following acceptance of this paper, we will publish the code for the analysis as well as the source data files for figures and tables online as part of an online repository that conforms to the FAIR (Findable, Accessible, Interoperable, and Re-usable) data principles. Additionally, the raw data are openly available from the HCP website which also conforms to the FAIR principles.

## 6 Acknowledgements

This work was supported by the René and Susanne Braginsky Foundation (KES), the Swiss National Science Foundation 320030_179377 (KES), and the University of Zurich (KES).

## Supplementary Tables

**Supplementary Table S1.**
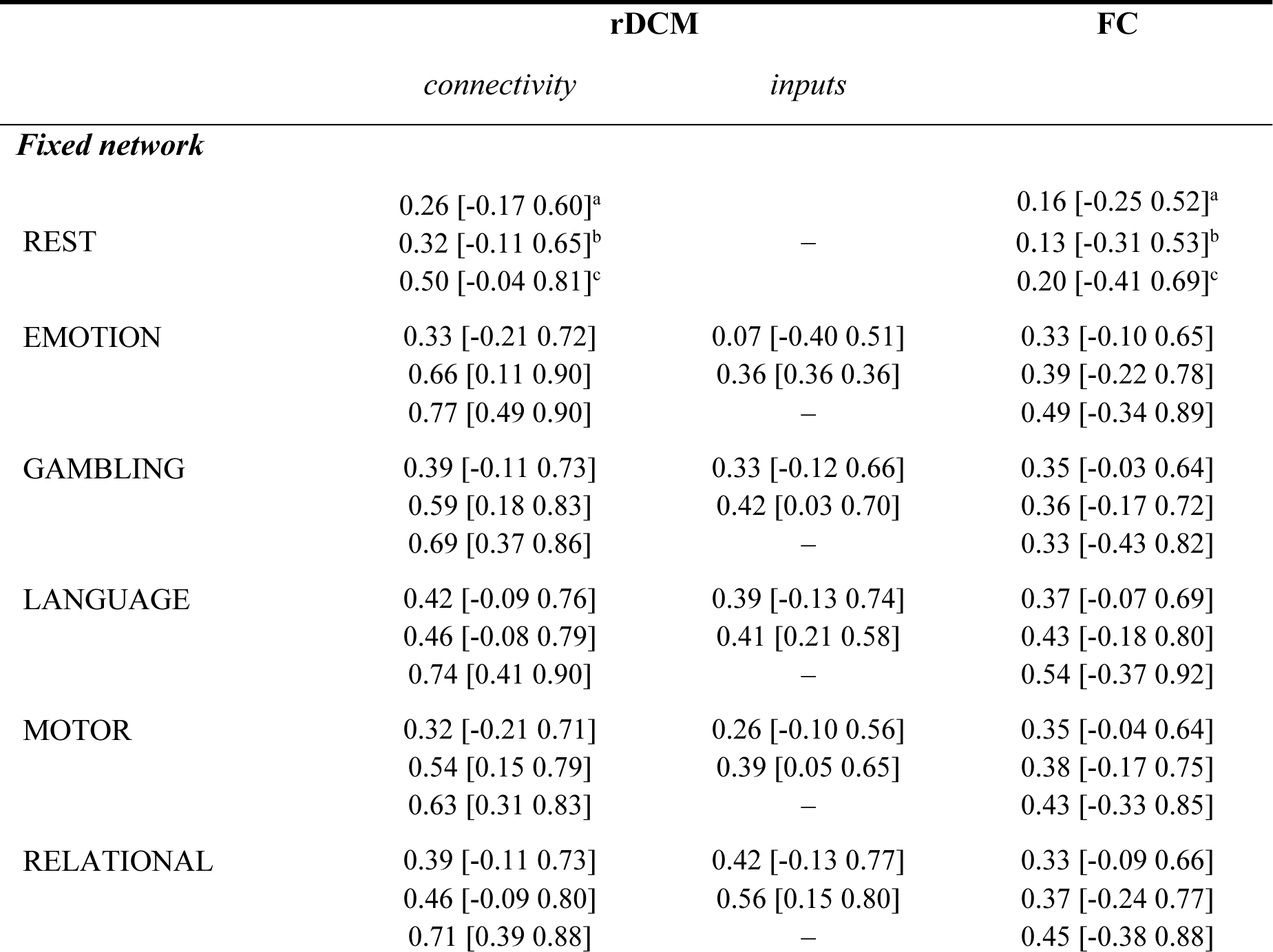
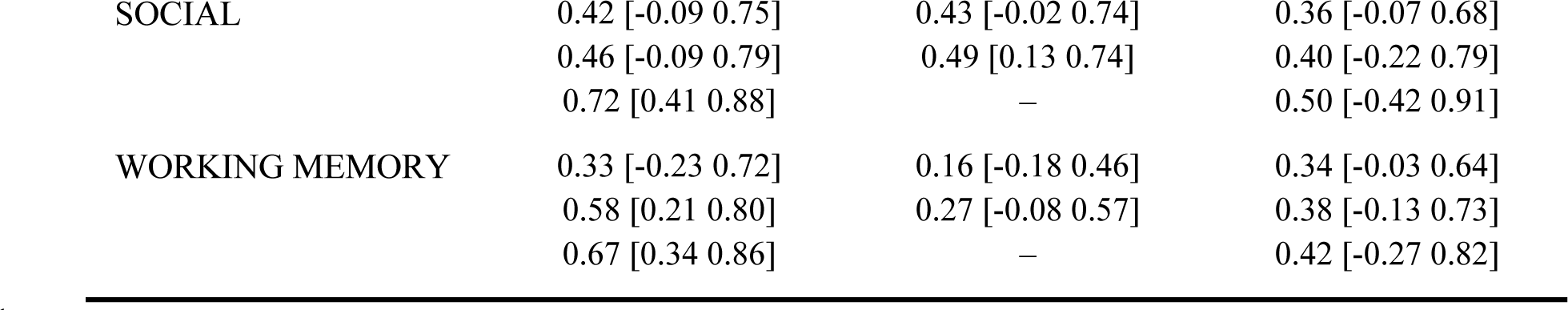
Test-retest reliability of model parameter estimates for regression DCM and functional connectivity. Test-retest reliability of parameter estimates was assessed in terms of the intra-class correlation coefficient (ICC) between estimates of session 1 (“test”) and session 2 (“retest”) for a fixed (full) network architecture (i.e., classical rDCM and Pearson’s correlation coefficient). Here, we report the mean (averaged across parameters) ICC value and 95% confidence interval (CI). Averaging of the connection-wise ICC values as well as computing the 95% CI was achieved by (i) transforming connection-wise ICC values to z-space using Fisher-z transformation, (ii) computing mean as well as lower and upper bound of the 95% CI in z-space, and finally (iii) back-transforming estimates to r-space. Test-retest reliability is reported for the connectivity and driving input parameters of rDCM (*middle*) as well as for the functional connectivity estimates (*right*). For both methods, results are shown for all HCP paradigms (i.e., rest, emotional processing, gambling, language, motor, relational processing, social cognition, and working memory) for the Schaefer atlas. Furthermore, results are shown for: (^a^) all parameters (*top row*), (^b^) significant parameters (*middle row*), and (^c^) top-1000 parameters (*bottom row*).

|  | rDCM |  | FC |
| --- | --- | --- | --- |
|  | *connectivity* | *inputs* |  |
| ***Fixed network*** |  |  |  |
| REST | 0.26 [-0.17 0.60]<sup>a</sup> | — | 0.16 [-0.25 0.52]<sup>a</sup> |
|  | 0.32 [-0.11 0.65]<sup>b</sup> |  | 0.13 [-0.31 0.53]<sup>b</sup> |
|  | 0.50 [-0.04 0.81]<sup>c</sup> |  | 0.20 [-0.41 0.69]<sup>c</sup> |
| EMOTION | 0.33 [-0.21 0.72] | 0.07 [-0.40 0.51] | 0.33 [-0.10 0.65] |
|  | 0.66 [0.11 0.90] | 0.36 [0.36 0.36] | 0.39 [-0.22 0.78] |
|  | 0.77 [0.49 0.90] | — | 0.49 [-0.34 0.89] |
| GAMBLING | 0.39 [-0.11 0.73] | 0.33 [-0.12 0.66] | 0.35 [-0.03 0.64] |
|  | 0.59 [0.18 0.83] | 0.42 [0.03 0.70] | 0.36 [-0.17 0.72] |
|  | 0.69 [0.37 0.86] | — | 0.33 [-0.43 0.82] |
| LANGUAGE | 0.42 [-0.09 0.76] | 0.39 [-0.13 0.74] | 0.37 [-0.07 0.69] |
|  | 0.46 [-0.08 0.79] | 0.41 [0.21 0.58] | 0.43 [-0.18 0.80] |
|  | 0.74 [0.41 0.90] | — | 0.54 [-0.37 0.92] |
| MOTOR | 0.32 [-0.21 0.71] | 0.26 [-0.10 0.56] | 0.35 [-0.04 0.64] |
|  | 0.54 [0.15 0.79] | 0.39 [0.05 0.65] | 0.38 [-0.17 0.75] |
|  | 0.63 [0.31 0.83] | — | 0.43 [-0.33 0.85] |
| RELATIONAL | 0.39 [-0.11 0.73] | 0.42 [-0.13 0.77] | 0.33 [-0.09 0.66] |
|  | 0.46 [-0.09 0.80] | 0.56 [0.15 0.80] | 0.37 [-0.24 0.77] |
|  | 0.71 [0.39 0.88] | — | 0.45 [-0.38 0.88] |
| SOCIAL | 0.42 [-0.09 0.75] | 0.43 [-0.02 0.74] | 0.36 [-0.07 0.68] |
|  | 0.46 [-0.09 0.79] | 0.49 [0.13 0.74] | 0.40 [-0.22 0.79] |
|  | 0.72 [0.41 0.88] | — | 0.50 [-0.42 0.91] |
| WORKING MEMORY | 0.33 [-0.23 0.72] | 0.16 [-0.18 0.46] | 0.34 [-0.03 0.64] |
|  | 0.58 [0.21 0.80] | 0.27 [-0.08 0.57] | 0.38 [-0.13 0.73] |
|  | 0.67 [0.34 0.86] | — | 0.42 [-0.27 0.82] |

**Supplementary Table S2.** Test-retest reliability of model parameter estimates for regression DCM and functional connectivity (sparsity constraints). Test-retest reliability of parameter estimates was assessed in terms of the intra-class correlation coefficient (ICC) between estimates of session 1 (“test”) and session 2 (“retest”) for sparsity constraints (i.e., rDCM with sparsity constraints and L1-regularized partial correlations). Here, we report the mean (averaged across parameters) ICC value and 95% confidence interval (CI). Averaging of the connection-wise ICC values as well as computing the 95% CI was done in z-space (see caption of Tab. 2 for details). Test-retest reliability is reported for the connectivity and driving input parameters of rDCM (*middle*) as well as for the functional connectivity estimates (*right*). For both methods, results are shown for all HCP paradigms (i.e., rest, emotional processing, gambling, language, motor, relational processing, social cognition, and working memory) for the Schaefer atlas. Furthermore, results are shown for: (^a^) all parameters (*top row*), (^b^) significant parameters (*middle row*), and (^c^) top-1000 parameters (*bottom row*).

|  | rDCM |  | FC |
| --- | --- | --- | --- |
|  | *connectivity* | *inputs* |  |
| ***Fixed network*** |  |  |  |
| REST | 0.03 [-0.27 0.32]<sup>a</sup> |  | 0.15 [-0.39 0.61]<sup>a</sup> |
|  | 0.03 [-0.26 0.31]<sup>b</sup> | – | 0.52 [0.04 0.80]<sup>b</sup> |
|  | 0.06 [-0.29 0.39]<sup>c</sup> |  | 0.57 [0.13 0.82]<sup>c</sup> |
| EMOTION | 0.25 [-0.24 0.64] | 0.19 [-0.26 0.57] | 0.05 [-0.41 0.49] |
|  | 0.46 [-0.16 0.82] | – | 0.33 [-0.01 0.60] |
|  | 0.70 [0.36 0.87] | – | 0.32 [-0.08 0.63] |
| GAMBLING | 0.30 [-0.16 0.64] | 0.27 [-0.19 0.63] | 0.06 [-0.44 0.53] |
|  | 0.33 [-0.12 0.66] | 0.41 [-0.03 0.71] | 0.30 [-0.08 0.60] |
|  | 0.60 [0.22 0.82] | – | 0.31 [-0.05 0.61] |
| LANGUAGE | 0.29 [-0.15 0.63] | 0.39 [-0.12 0.74] | 0.09 [-0.44 0.57] |
|  | 0.31 [-0.09 0.63] | 0.38 [0.15 0.57] | 0.39 [-0.03 0.70] |
|  | 0.56 [0.11 0.82] | – | 0.40 [-0.02 0.70] |
| MOTOR | 0.35 [-0.10 0.68] | 0.19 [-0.35 0.64] | 0.07 [-0.39 0.50] |
|  | 0.43 [0.05 0.70] | – | 0.33 [-0.05 0.63] |
|  | 0.49 [0.09 0.75] | – | 0.35 [-0.05 0.65] |
| RELATIONAL | 0.31 [-0.13 0.65] | 0.41 [-0.13 0.76] | 0.08 [-0.43 0.54] |
|  | 0.34 [-0.07 0.65] | 0.53 [0.09 0.80] | 0.35 [0.00 0.63] |
|  | 0.60 [0.19 0.83] | – | 0.35 [-0.04 0.64] |
| SOCIAL | 0.32 [-0.12 0.65] | 0.38 [-0.08 0.70] | 0.09 [-0.46 0.59] |
|  | 0.33 [-0.09 0.64] | 0.45 [0.10 0.71] | 0.39 [-0.01 0.69] |
|  | 0.59 [0.16 0.83] | – | 0.42 [0.01 0.71] |
| WORKING MEMORY | 0.27 [-0.14 0.60] | 0.12 [-0.22 0.44] | 0.08 [-0.38 0.50] |
|  | 0.27 [-0.08 0.56] | – | 0.35 [-0.04 0.64] |
|  | 0.30 [-0.10 0.62] | – | 0.36 [-0.02 0.65] |

## Supplementary Figures

**Supplementary Figure S1:**
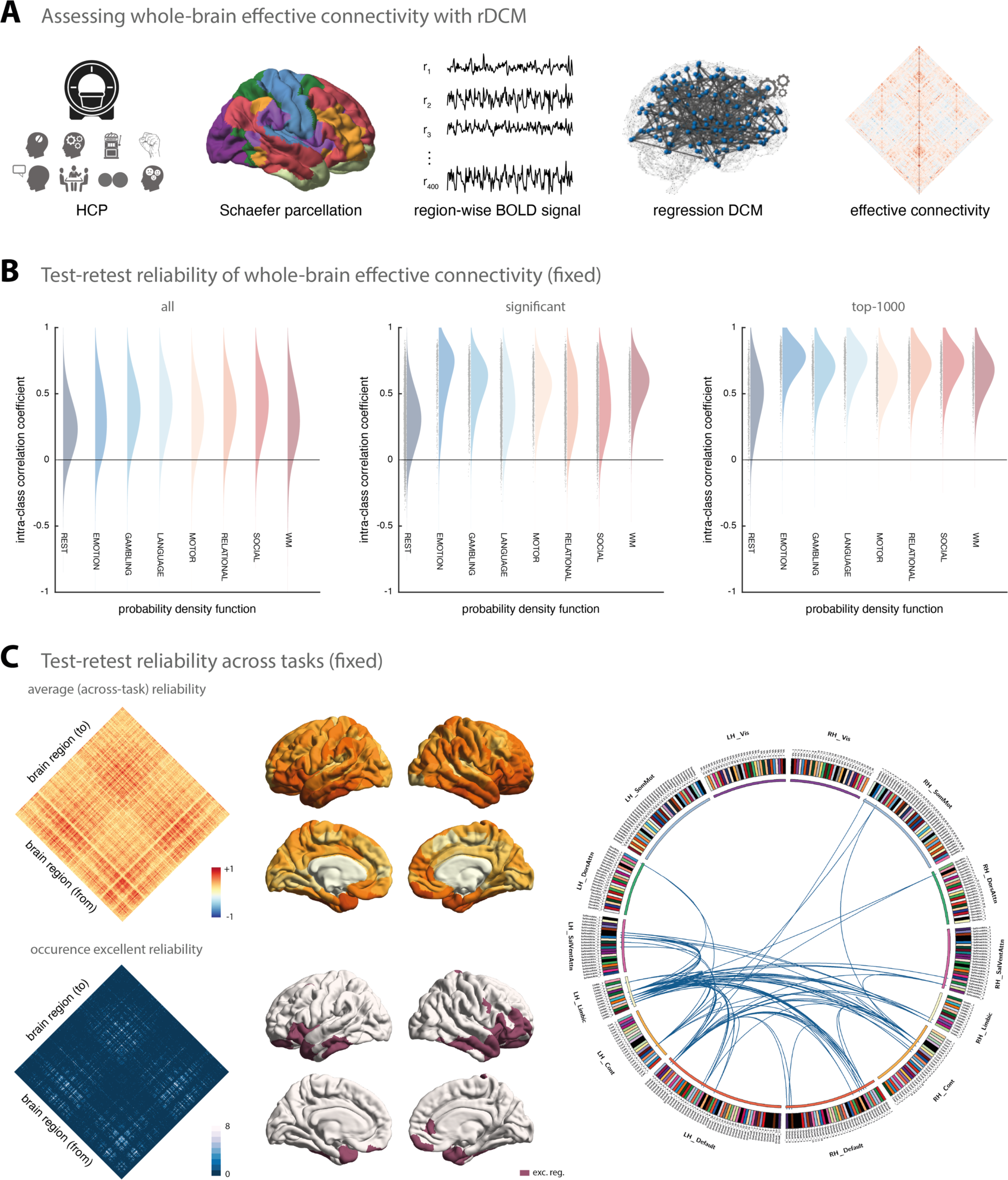
Test-retest reliability of regression DCM for a fixed network architecture. **(A)** Methodological overview. Resting-state and task-based fMRI data from the Human Connectome Project (HCP) is used for the analysis. Region-wise BOLD signal time series were extracted from a whole-brain parcellation scheme (e.g., the Glasser atlas) and effective connectivity was inferred at the whole-brain level using regression dynamic causal modeling (rDCM). The rDCM parameter estimates were then subjected to group-level consistency and test-retest reliability analyses. **(B)** Estimates of the probability density functions (using the nonparametric kernel smoothing of *fitdist.m* implemented in MATLAB) of the connection-wise intra-class correlation coefficient (ICC) for the resting-state and all 7 tasks (i.e., emotional processing, gambling, language, motor, relational processing, social cognition, and working memory) for the Schaefer atlas. Results are shown when considering all connections (*left*), significant connections (*middle*), and the top-1000 connections (*right*). **(C)** Mean (averaged across all paradigms) test-retest reliability (*top, left*) as well as how often (i.e., in how many paradigms) a connection showed excellent reliability (*bottom, left*). Mean test-retest reliability projected onto the cortical surface (*top, middle*) and the cortical location of all regions that are linked via connections that show excellent reliability in all 8 paradigms (*bottom, middle*). Connectogram showing the connections with excellent reliability in all 8 paradigms (*right*). The connectogram was produced using Circos (publicly available at http://circos.ca/software/). Parcels of the Schaefer atlas can be assigned to the resting state networks (RSN) defined in Yeo et al. (2011), which include visual (Vis), somatosensory-motor (SomMot), dorsal attention (DorsAttn), ventral attention (SalVentAttn), limbic (Limbic), control (Cont), and default mode network (Default).

**Supplementary Figure S2:**
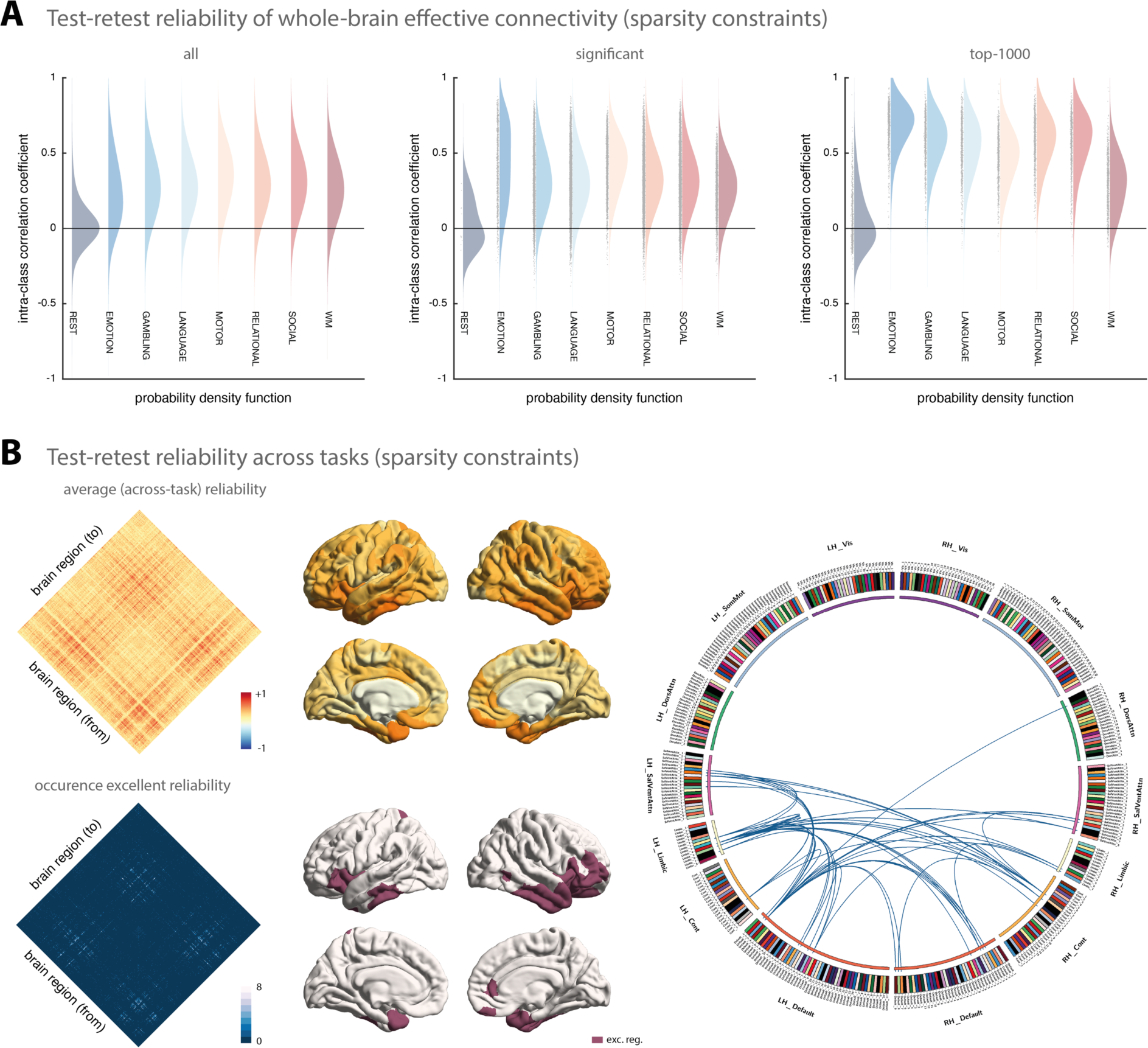
Test-retest reliability of regression DCM with sparsity constraints. **(A)** Estimates of the probability density functions (using the nonparametric kernel smoothing of *fitdist.m* implemented in MATLAB) of the connection-wise intra-class correlation coefficient (ICC) for the resting-state and all 7 tasks (i.e., emotional processing, gambling, language, motor, relational processing, social cognition, and working memory) for the Schaefer atlas. Results are shown when considering all connections (*left*), significant connections (*middle*), and the top-1000 connections (*right*). **(B)** Mean (averaged across all paradigms) test-retest reliability (*top, left*) as well as how often (i.e., in how many paradigms) a connection showed excellent reliability (*bottom, left*). Mean test-retest reliability projected onto the cortical surface (*top, middle*) and the cortical location of all regions that are linked via connections that show excellent reliability in at least 6 paradigms (*bottom, middle*). Connectogram showing the connections with excellent reliability in at least 6 paradigms (*right*). The connectogram was produced using Circos (publicly available at http://circos.ca/software/). Parcels of the Schaefer atlas can be assigned to the resting state networks (RSN) defined in Yeo et al. (2011), which include visual (Vis), somatosensory-motor (SomMot), dorsal attention (DorsAttn), ventral attention (SalVentAttn), limbic (Limbic), control (Cont), and default mode network (Default).

**Supplementary Figure S3:**
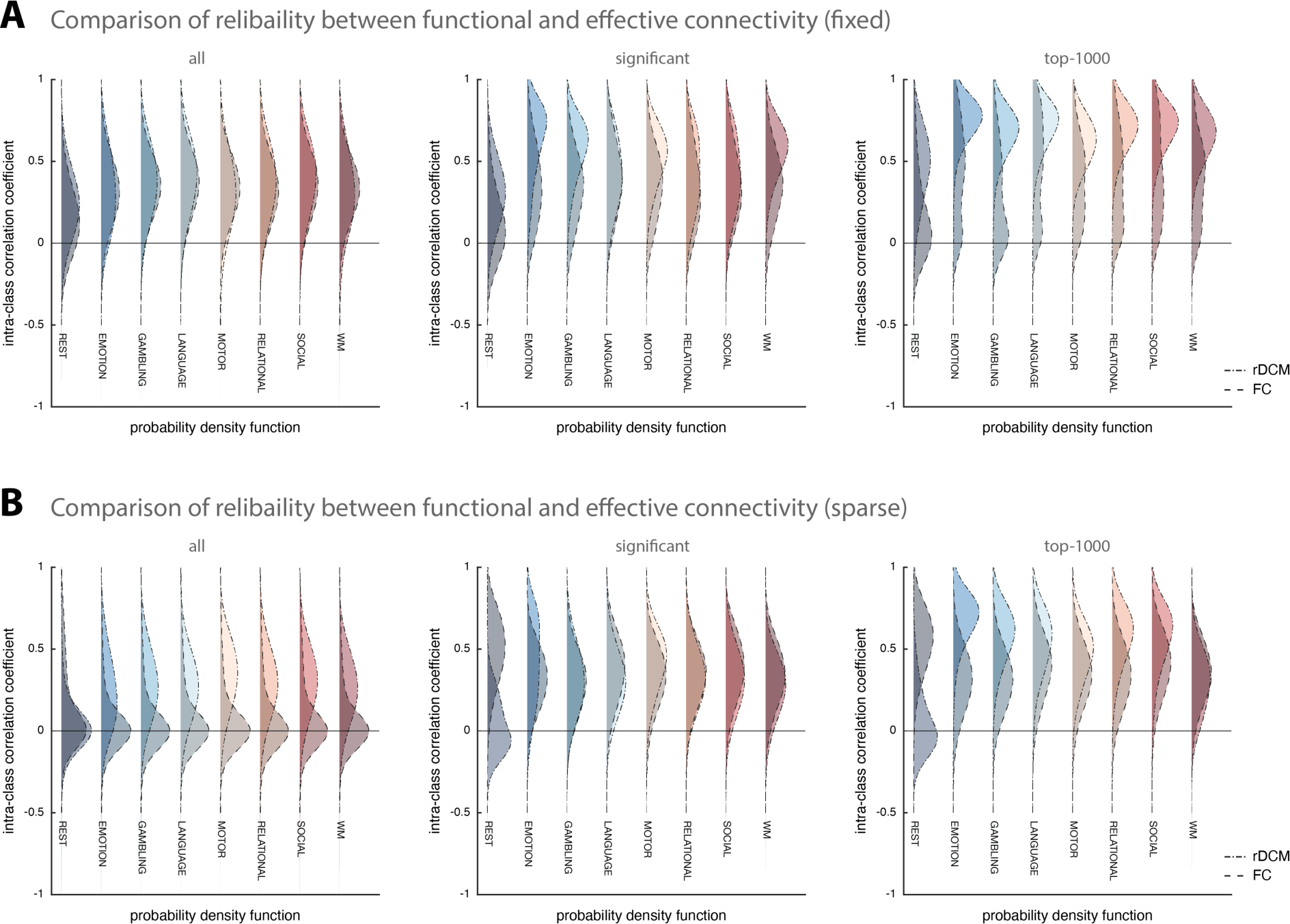
Comparison of test-retest reliability between regression DCM and functional connectivity. **(A)** Estimates of the probability density functions (using the nonparametric kernel smoothing of *fitdist.m* implemented in MATLAB) of the connection-wise intra-class correlation coefficient (ICC) for the resting-state and all 7 tasks (i.e., emotional processing, gambling, language, motor, relational processing, social cognition, and working memory) for the Schaefer atlas for fixed (full) connectivity methods (i.e., classical rDCM and Pearson’s correlation coefficient), and **(B)** sparse connectivity methods (i.e., rDCM with sparsity constraints and L1-regularized partial correlations). Probability density functions representing rDCM results are shown with dot-dashed lines and lighter colors, whereas probability density functions representing functional connectivity results are shown with dashed lines and darker colors. For each connectivity variant, results are shown when considering all connections (*left*), significant connections (*middle*), and the top-1000 connections (*right*).

## Footnotes

1 The number of driving input parameters varied for the different tasks because a different number of driving input regressors was available for each task.

